# Shared heritability and functional enrichment across six solid cancers

**DOI:** 10.1101/453480

**Authors:** Xia Jiang, Hilary K. Finucane, Fredrick R. Schumacher, Stephanie L. Schmit, Jonathan P. Tyrer, Younghun Han, Kyriaki Michailidou, Corina Lesseur, Karoline B. Kuchenbaecker, Joe Dennis, David V. Conti, Graham Casey, Mia M. Gaudet, Jeroen R. Huyghe, Demetrius Albanes, Melinda C. Aldrich, Angeline S. Andrew, Irene L. Andrulis, Hoda Anton-Culver, Antonis C. Antoniou, Natalia N. Antonenkova, Susanne M. Arnold, Kristan J. Aronson, Banu K. Arun, Elisa V. Bandera, Rosa B. Barkardottir, Daniel R. Barnes, Jyotsna Batra, Matthias W. Beckmann, Javier Benitez, Sara Benlloch, Andrew Berchuck, Sonja I. Berndt, Heike Bickeböller, Stephanie A. Bien, Carl Blomqvist, Stefania Boccia, Natalia V. Bogdanova, Stig E. Bojesen, Manjeet K. Bolla, Hiltrud Brauch, Hermann Brenner, James D. Brenton, Mark N. Brook, Joan Brunet, Hans Brunnström, Daniel D. Buchanan, Barbara Burwinkel, Ralf Butzow, Gabriella Cadoni, Trinidad Caldés, Maria A. Caligo, Ian Campbell, Peter T. Campbell, Géraldine Cancel-Tassin, Lisa Cannon-Albright, Daniele Campa, Neil Caporaso, André L. Carvalho, Andrew T. Chan, Jenny Chang-Claude, Stephen J. Chanock, Chu Chen, David C. Christiani, Kathleen B.M. Claes, Frank Claessens, Judith Clements, J. Margriet Collée, Marcia Cruz Correa, Fergus J. Couch, Angela Cox, Julie M. Cunningham, Cezary Cybulski, Kamila Czene, Mary B. Daly, Anna deFazio, Peter Devilee, Orland Diez, Manuela Gago-Dominguez, Jenny L. Donovan, Thilo Dörk, Eric J. Duell, Alison M. Dunning, Miriam Dwek, Diana M. Eccles, Christopher K. Edlund, Digna R Velez Edwards, Carolina Ellberg, D. Gareth Evans, Peter A. Fasching, Robert L. Ferris, Triantafillos Liloglou, Jane C. Figueiredo, Olivia Fletcher, Renée T. Fortner, Florentia Fostira, Silvia Franceschi, Eitan Friedman, Steven J. Gallinger, Patricia A. Ganz, Judy Garber, José A. García-Sáenz, Simon A. Gayther, Graham G. Giles, Andrew K. Godwin, Mark S. Goldberg, David E. Goldgar, Ellen L. Goode, Marc T. Goodman, Gary Goodman, Kjell Grankvist, Mark H. Greene, Henrik Gronberg, Jacek Gronwald, Pascal Guénel, Niclas Håkansson, Per Hall, Ute Hamann, Freddie C. Hamdy, Robert J. Hamilton, Jochen Hampe, Aage Haugen, Florian Heitz, Rolando Herrero, Peter Hillemanns, Michael Hoffmeister, Estrid Høgdall, Chul Yun-Hong, John L. Hopper, Richard Houlston, Peter J. Hulick, David J. Hunter, David G. Huntsman, Gregory Idos, Evgeny N. Imyanitov, Sue Ann Ingles, Claudine Isaacs, Anna Jakubowska, Paul James, Mark A. Jenkins, Mattias Johansson, Mikael Johansson, Esther M. John, Amit D. Joshi, Radka Kaneva, Beth Y. Karlan, Linda E. Kelemen, Tabea Kühl, Tee Kay-Khaw, Elza Khusnutdinova, Adam S. Kibel, Lambertus A. Kiemeney, Jeri Kim, Susanne K. Kjaer, Julia A. Knight, Manolis Kogevinas, ZSofia Kote-Jarai, Stella Koutros, Vessela N. Kristensen, Jolanta Kupryjanczyk, Martin Lacko, Stephan Lam, Diether Lambrechts, Maria Teresa Landi, Philip Lazarus, Nhu D. Le, Eunjung Lee, Flavio Lejbkowicz, Josef Heinz-Lenz, Goska Leslie, Davor Lessel, Jenny Lester, Douglas A. Levine, Li Li, Christopher I. Li, Annika Lindblom, Noralane M. Lindor, Geoffrey Liu, Fotios Loupakis, Jan Lubinński, Lovise Maehle, Christiane Maier, Arto Mannermaa, Loic Le Marchand, Sara Margolin, Taymaa May, Lesley McGuffog, Alfons Meindl, Pooja Middha, Austin Miller, Roger L. Milne, Robert J. MacInnis, Francesmary Modugno, Marco Montagna, Victor Moreno, Kirsten B. Moysich, Lorelei Mucci, Kenneth Muir, Anna Marie Mulligan, Katherine L. Nathanson, David E. Neal, Andrew R. Ness, Susan L. Neuhausen, Heli Nevanlinna, Polly A. Newcomb, Lisa F. Newcomb, Finn Cilius Nielsen, Liene Nikitina-Zake, Børge G. Nordestgaard, Robert L. Nussbaum, Kenneth Offit, Edith Olah, Ali Amin Al Olama, Olufunmilayo I. Olopade, Andrew F. Olshan, Håkan Olsson, Ana Osorio, Hardev Pandha, Jong Y. Park, Nora Pashayan, Michael T. Parsons, Tanja Pejovic, Kathryn L. Penney, Wilbert HM. Peters, Catherine M. Phelan, Amanda I. Phipps, Dijana Plaseska-Karanfilska, Miranda Pring, Darya Prokofyeva, Paolo Radice, Kari Stefansson, Susan J. Ramus, Leon Raskin, Gad Rennert, Hedy S. Rennert, Elizabeth J. van Rensburg, Marjorie J. Riggan, Harvey A. Risch, Angela Risch, Monique J. Roobol, Barry S. Rosenstein, Mary Anne Rossing, Kim De Ruyck, Emmanouil Saloustros, Dale P. Sandler, Elinor J. Sawyer, Matthew B. Schabath, Johanna Schleutker, Marjanka K. Schmidt, V. Wendy Setiawan, Hongbing Shen, Erin M. Siegel, Weiva Sieh, Christian F. Singer, Martha L. Slattery, Karina Dalsgaard Sorensen, Melissa C. Southey, Amanda B. Spurdle, Janet L. Stanford, Victoria L. Stevens, Sebastian Stintzing, Jennifer Stone, Karin Sundfeldt, Rebecca Sutphen, Anthony J. Swerdlow, Eloiza H. Tajara, Catherine M. Tangen, Adonina Tardon, Jack A. Taylor, M. Dawn Teare, Manuel R. Teixeira, Mary Beth Terry, Kathryn L. Terry, Stephen N. Thibodeau, Mads Thomassen, Line Bjørge, Marc Tischkowitz, Amanda E. Toland, Diana Torres, Paul A. Townsend, Ruth C. Travis, Nadine Tung, Shelley S. Tworoger, Cornelia M. Ulrich, Nawaid Usmani, Celine M. Vachon, Els Van Nieuwenhuysen, Ana Vega, Elías Miguel Aguado-Barrera, Qin Wang, Penelope M. Webb, Clarice R. Weinberg, Stephanie Weinstein, Mark C. Weissler, Jeffrey N. Weitzel, Catharine ML West, Emily White, Alice S. Whittemore, Erich H-Wichmann, Fredrik Wiklund, Robert Winqvist, Alicja Wolk, Penella Woll, Michael Woods, Anna H. Wu, Xifeng Wu, Drakoulis Yannoukakos, Wei Zheng, Shanbeh Zienolddiny, Argyrios Ziogas, Kristin K. Zorn, Jacqueline M. Lane, Richa Saxena, Duncan Thomas, Rayjean J. Hung, Brenda Diergaarde, James McKay, Ulrike Peters, Li Hsu, Montserrat García-Closas, Rosalind A. Eeles, Georgia Chenevix-Trench, Paul J. Brennan, Christopher A. Haiman, Jacques Simard, Douglas F. Easton, Stephen B. Gruber, Paul D.P. Pharoah, Alkes L. Price, Bogdan Pasaniuc, Christopher I. Amos, Peter Kraft, Sara Lindström, BCAC, CCFR, CIMBA, CORECT, GECCO, OCAC, PRACTICAL, CRUK, BPC3, CAPS, PEGASUS, TRICL-ILCCO, ABCTB, APCB BioResource, BCFR, CONSIT TEAM, EMBRACE, GC-HBOC, GEMO, HEBON, kConFab/AOCS Mod SQuaD and SWE-BRCA

**Affiliations:** Program in Genetic Epidemiology and Statistical Genetics, Harvard T.H. Chan School of Public Health, 677 Huntington Ave, Boston, MA, 02115, USA.; Unit of Cardiovascular Epidemiology, Institute of Environmental Medicine, Karolinska Institutet. Nobels vagen 13, Stockholm, 17177, Sweden.; Department of Epidemiology, Harvard T.H. Chan School of Public Health, 677 Huntington Ave, Boston, MA, 02115, USA.; Program in Medical and Population Genetics, Broad Institute of MIT and Harvard, 75 Ames St, Cambridge, MA, 02142, USA.; Department of Population and Quantitative Health Sciences, Case Western Reserve University, 10900 Eucid Avenue, Cleveland, OH, 44106, USA.; Seidman Cancer Center, University Hospitals, Cleveland, OH, 44106, USA.; Department of Cancer Epidemiology, H. Lee Moffitt Cancer Center and Research Institute, 12902 Magnolia Dr. MRC-CANCONT, Tampa, FL, 33612, USA.; Department of Gastrointestinal Oncology, H. Lee Moffitt Cancer Center and Research Institute, 12902 Magnolia Dr. MRC-CANCONT, Tampa, FL, 33612, USA.; Centre for Cancer Genetic Epidemiology, Department of Oncology, University of Cambridge, 2 Worts’ Causeway, Cambridge, CB1 8RN, UK.; Department of Biomedical Data Science, The Geisel School of Medicine at Dartmouth, 1 Medical Center Drive, Lebanon, NH, 03756, USA.; Centre for Cancer Genetic Epidemiology, Department of Public Health and Primary Care, University of Cambridge, 2 Worts’ Causeway, Cambridge, CB1 8RN, UK.; Department of Electron Microscopy/Molecular Pathology, The Cyprus Institute of Neurology and Genetics, 1683 Nicosia, Nicosia, Cyprus.; Genetic Epidemiology Group, International Agency for Research on Cancer, 150 Cours Albert Thomas, Lyon, 69008, France.; Section of Genetics, International Agency for Research on Cancer, 150 cours Albert Thomas, Lyon, 69008, France.; Division of Psychiatry, University College London, Maple House, 149 Tottenham Court Road, London, W1T 7NF, UK.; UCL Genetics Institute, University College London, Gower Street, London, WC1E 6BT, UK.; Department of Preventive Medicine, Keck School of Medicine, University of Southern California Norris Comprehensive Cancer Center, Los Angeles, CA, 48109, USA.; Public Health Sciences, University of Virginia, P.O. Box 800717, Charlottesville, VI, 22908, USA.; Center for Public Health Genomics, University of Virginia, P.O. Box 800717, Charlottesville, VI, 22908, USA.; Epidemiology Research Program, American Cancer Society, 250 Williams Street NW, Atlanta, GA, 30303, USA.; Public Health Sciences Division, Fred Hutchinson Cancer Research Center, 1100 Fairview Ave. N., Seattle, WA 98109-1024, USA.; Division of Cancer Epidemiology and Genetics, National Cancer Institute, 9609 Medical Center Dr, Rockville, MD, 20850, USA.; Department of Thoracic Surgery, Division of Epidemiology, Vanderbilt University Medical Center, 609 Oxford House, Nashville, TN 37232, USA.; Department of Neurology, Dartmouth-Hitchcock Medical Center, 7927 Rubin Building, Room 860, One Medical Center Drive, Lebanon, NH, 3756, USA.; Fred A. Litwin Center for Cancer Genetics, Lunenfeld-Tanenbaum Research Institute of Mount Sinai Hospital, 600 University Avenue, Toronto, ON, M5G1X5, Canada.; Department of Molecular Genetics, University of Toronto, 1 King’s College Circle, Toronto, ON, M5S1A8, Canada.; Department of Epidemiology, Genetic Epidemiology Research Institute, University of California Irvine, 224 Irvine Hall, Irvine, CA, 92617, USA.; N.N. Alexandrov Research Institute of Oncology and Medical Radiology, Settlement of Lesnoy-2, Minsk, 223040, Belarus.; Markey Cancer Center, University of Kentucky, 800 Rose Street, cc445, Lexington, KY, 40508, USA.; Department of Public Health Sciences, and Cancer Research Institute, Queen’s University, 10 Stuart Street, Kingston, ON, K7L 3N6, Canada.; Department of Breast Medical Oncology, University of Texas MD Anderson Cancer Center, 1155 Pressler St, Houston, TX, 77030, USA.; Cancer Prevention and Control Program, Rutgers Cancer Institute of New Jersey, 195 Little Albany Street, Room 5568, New Brunswick, NJ, 08903, USA.; Department of Pathology, Landspitali University Hospital, Hringbraut, Reykjavik, 101, Iceland.; BMC (Biomedical Centre), Faculty of Medicine, University of Iceland, Vatnsmyrarvegi 16, Reykjavik, 101, Iceland.; Australian Prostate Cancer Research Centre-Qld, Translational Research Institute, 37 Kent St, Woolloongabba, Queensland, 4102, Australia.; Institute of Health and Biomedical Innovation and School of Biomedical Science, Queensland University of Technology, 60 Musk Ave, Kelvin Grove, Queensland, 4059, Australia.; Department of Gynecology and Obstetrics, Comprehensive Cancer Center Erlangen Nuremberg, University Hospital Erlangen, Friedrich-Alexander-University Erlangen-Nuremberg, Universitaetsstrasse 21-23, Erlangen, 91054, Germany.; Human Cancer Genetics Programme, Spanish National Cancer Research Centre (CNIO), Calle de Melchor Fernández Almagro, 3, Madrid 28029, Spain.; Biomedical Network on Rare Diseases (CIBERER), Av. Monforte de Lemos, 3-5. Pabellón 11. Planta 0, Madrid, 28029, Spain.; Division of Genetics and Epidemiology, The Institute of Cancer Research, London, SM2 5NG, UK.; Department of Obstetrics and Gynecology, Duke University Medical Center, 25171 Morris Bldg, Durham, NC, 27710, USA.; Department of Genetic Epidemiology, University Medical Center Goettingen, Humboldtallee 32, Goettingen, 37073, Germany.; School of Public Health, University of Washington, 1959 NE Pacific Street, Health Science Buidling, F-350, Seattle WA 98195, USA.; Department of Oncology, Helsinki University Hospital, University of Helsinki, Haartmaninkatu 4, Helsinki, 00290, Finland.; Department of Oncology, örebro University Hospital, örebro, 70185, Sweden.; Fondazione Policlinico Universitario A. Gemelli IRCCS, Roma, 00168, Italia.; Università Cattolica del Sacro Cuore, Roma, 00168, Italia.; Department of Radiation Oncology, Hannover Medical School, Carl-Neuberg-Straße 1, Hannover, 30625, Germany.; Gynaecology Research Unit, Hannover Medical School, Carl-Neuberg-Straße 1, Hannover, 30625, Germany.; Copenhagen General Population Study, Herlev and Gentofte Hospital, Copenhagen University Hospital, Herlev Ringvej 75, Herlev, 2730, Denmark.; Department of Clinical Biochemistry, Herlev and Gentofte Hospital, Copenhagen University Hospital, Herlev Ringvej 75, Herlev, 2730, Denmark.; Faculty of Health and Medical Sciences, University of Copenhagen, Blegdamsvej 3B, Copenhagen, 2200, Denmark.; Dr. Margarete Fischer-Bosch-Institute of Clinical Pharmacology, Auerbachstr. 112, Stuttgart, 70376, Germany.; University of Tübingen, Geschwister-Scholl-Platz, Tübingen, 72074, Germany.; German Cancer Consortium (DKTK), German Cancer Research Center (DKFZ), Im Neuenheimer Feld 280, Heidelberg, 69120, Germany.; Division of Clinical Epidemiology and Aging Research, German Cancer Research Center (DKFZ), Im Neuenheimer Feld 280, Heidelberg, 69120, Germany.; Division of Preventive Oncology, German Cancer Research Center (DKFZ) and National Center for Tumor Diseases (NCT), Im Neuenheimer Feld 280, Heidelberg, 69120, Germany.; Cancer Research UK Cambridge Institute, University of Cambridge, Li Ka Shing Centre, Robinson Way, Cambridge, CB2 0RE, UK.; Genetic Counseling Unit, Hereditary Cancer Program, IDIBGI (Institut d’Investigació Biomèdica de Girona), Catalan Institute of Oncology, CIBERONC, Av. França s/n., Girona, 17007, Spain.; Clinical Sciences, Lund University, Box 117, Lund, 221 00, Sweden.; Department of Genetics and Pathology, Division of Laboratory Medicine, 221 85 Lund, Sweden.; University of Melbourne Centre for Cancer Research, Victorian Comprehensive Cancer Centre, Parkville, Victoria, 3010, Australia.; Colorectal Oncogenomics Group, Department of Clinical Pathology, The University of Melbourne, Parkville, Victoria 3010 Australia.; Genomic Medicine and Family Cancer Clinic, Royal Melbourne Hospital, Parkville, Victoria 3010 Australia.; Department of Obstetrics and Gynecology, University of Heidelberg, Im Neuenheimer Feld 440, Heidelberg, 69120, Germany.; Molecular Epidemiology Group, C080, German Cancer Research Center (DKFZ), Im Neuenheimer Feld 280, Heidelberg, 69120, Germany.; Department of Pathology, University of Helsinki and Helsinki University Hospital, Biomedicum Helsinki 4th floor, Haartmaninkatu 8, Helsinki, 00029, Finland.; Medical Oncology Department, Hospital Clínico San Carlos, Instituto de Investigación Sanitaria San Carlos (IdISSC), Centro Investigación Biomédica en Red de Cáncer (CIBERONC), Calle del Prof Martín Lagos, Madrid, 28040, Spain.; Section of Genetic Oncology, Dept. of Laboratory Medicine, University and University Hospital of Pisa, via Roma 67, Pisa, 56126, Italy.; Peter MacCallum Cancer Center, 305 Grattan Street, Melbourne, Victoria, 3000, Australia.; Sir Peter MacCallum Department of Oncology, The University of Melbourne, 305 Grattan Street, Melbourne, Victoria, 3000, Australia.; Sorbonne Université, GRC N°5 ONCOTYPE-URO, Tenon Hospital, Paris, France.; CeRePP, Tenon Hospital, Paris, France.; Division of Genetic Epidemiology, Department of Medicine, University of Utah School of Medicine, Salt Lake City, Utah, 84112, USA.; George E. Wahlen Department of Veterans Affairs Medical Center, Salt Lake City, Utah, USA.; Division of Cancer Epidemiology, German Cancer Research Center (DKFZ), Im Neuenheimer Feld 280, Heidelberg, 69120, Germany.; Department of Biology, University of Pisa, Pisa, 56126 Italy.; Molecular Oncology Research Center, Barretos Cancer Hospital, Rua Antenor Duarte Villela, 1331, São Paulo, 784-400, Brazil.; Head and Neck Surgery Department, Barretos Cancer Hospital, Pio XII, 1331, Antenor Duarte Villela St, Barretos, SP, 14784-400, Brazil.; Division of Gastroenterology, Massachusetts General Hospital, 55 Fruit Street, Boston, MA 02114, USA.; Channing Division of Network Medicine, Department of Medicine, Brigham and Women’s Hospital, Harvard Medical School, 181 Longwood Avenue, Boston, MA, 02115, USA.; Cancer Epidemiology Group, University Cancer Center Hamburg (UCCH), University Medical Center Hamburg-Eppendorf, Martinistraße 52, Hamburg, 20246, Germany.; Program in Epidemiology, Division of Public Health Sciences, Fred Hutchinson Cancer Research Center, 1100 Fairview Ave N, Seattle, WA, 98109, USA.; Centre for Medical Genetics, Ghent University, De Pintelaan 185, Gent, 9000, Belgium.; Molecular Endocrinology Laboratory, Department of Cellular and Molecular Medicine, KU Leuven, Leuven, Belgium.; Department of Clinical Genetics, Erasmus University Medical Center, Wytemaweg 80, Rotterdam, 3015 CN, The Netherlands.; University of Puerto Rico Medical Sciences Campus and Comprehensive Cancer Center, San Juan 00936 Puerto Rico.; Department of Laboratory Medicine and Pathology, Mayo Clinic, 200 First St. SW, Rochester, MN, 55905, USA.; Sheffield Institute for Nucleic Acids (SInFoNiA), Department of Oncology and Metabolism, University of Sheffield, Western Bank, Sheffield, S10 2TN, UK.; International Hereditary Cancer Center, Department of Genetics and Pathology, Pomeranian Medical University, ul. Unii Lubelskiej 1, 71-252 Szczecin, Poland.; Department of Medical Epidemiology and Biostatistics, Karolinska Institutet, Karolinska Univ Hospital, Stockholm, 171 76, Sweden.; Department of Clinical Genetics, Fox Chase Cancer Center, 333 Cottman Ave, Philadelphia, PA, 19111, USA.; Centre for Cancer Research, The Westmead Institute for Medical Research, The University of Sydney, 176 Hawkesbury Rd, Sydney, New South Wales, 2145, Australia.; Department of Gynaecological Oncology, Westmead Hospital, Hawkesbury Rd & Darcy Rd, Sydney, New South Wales, 2145, Australia.; Department of Pathology, Leiden University Medical Center, Albinusdreef 2, Leiden, 2333 ZA, TheNetherlands.; Department of Human Genetics, Leiden University Medical Center, Albinusdreef 2, Leiden, 2333 ZA, The Netherlands.; Oncogenetics Group, Clinical and Molecular Genetics Area, Vall d’Hebron Institute of Oncology (VHIO), University Hospital, Vall d’Hebron, Passeig de la Vall d’Hebron 119-129, Barcelona, 08035, Spain.; Genomic Medicine Group, Galician Foundation of Genomic Medicine, Instituto de Investigación Sanitaria de Santiago de Compostela (IDIS), Complejo Hospitalario Universitario de Santiago, SERGAS, Travesía da Choupana S/N, Santiago de Compostela, 15706, Spain.; Moores Cancer Center, University of California San Diego, 3855 Health Sciences Drive, La Jolla, CA, 92037, USA.; School of Social and Community Medicine, University of Bristol, Bristol, BS8 1TH, UK.; Unit of Nutrition and Cancer, Cancer Epidemiology Research Program, Catalan Institute of Oncology (ICO-IDIBELL), Av. Gran Via 199-203, L’Hospitalet de Llobregat, 08908 Barcelona, Spain.; Department of Biomedical Sciences, Faculty of Science and Technology, University of Westminster, 309 Regent Street, London, W1B 2HW, UK.; Cancer Sciences Academic Unit, Faculty of Medicine, University of Southampton, University Hospital Southampton, Tremona Road, SO16 6YD, UK.; Department of Medicine, Keck School of Medicine, University of Southern California, Los Angeles, CA, 90033, USA.; Vanderbilt Epidemiology Center, Vanderbilt Genetics Institute, Department of Obstetrics and Gynecology, Vanderbilt University Medical Center, 2525 West End Avenue, Suite 600, Nashville, TN, 37203, USA.; Department of Cancer Epidemiology, Clinical Sciences, Lund University, Barngatan 4, Skånes universitetssjukhus, Lund, 222 42, Sweden.; Manchester Centre for Genomic Medicine, Division of Evolution and Genomic Sciences, University of Manchester, St Mary’s Hospital, Central Manchester University Hospitals NHS Foundation Trust, Oxford Road, Manchester, M13 9WL, UK.; David Geffen School of Medicine, Department of Medicine Division of Hematology and Oncology, University of California at Los Angeles, 10833 Le Conte Ave, Los Angeles, CA, 90095, USA.; Department of Otolaryngology, University of Pittsburgh, UPMC Hillman Cancer Center, Cancer Pavilion, Suite 500, 5150 Centre Avenue, Pittsburgh, PA, 15232, USA.; Molecular and Clinical Cancer Medicine, Roy Castle Lung Cancer Research Programme, The University of Liverpool Institute of Translational Medicine, The Wiliam Duncan Building, 6 West Derby Street, Liverpool, L69 3BX, UK.; Samuel Oschin Comprehensive Cancer Institute, Cedars-Sinai Medical Center, 8700 Beverly Boulevard, Los Angeles, CA, 90048, USA.; Keck School of Medicine, University of Southern California, 1450 Biggy Street, Los Angeles, CA, 90033, USA.; The Breast Cancer Now Toby Robins Research Centre, The Institute of Cancer Research, 123 Old Brompton Road, London, SW7 3RP, UK.; Molecular Diagnostics Laboratory, INRASTES, National Centre for Scientific Research ‘Demokritos’, Neapoleos 10, Ag. Paraskevi, Athens, 15310, Greece.; Section of Infections, International Agency for Research on Cancer, 150 cours Albert Thomas, Lyon, 69008, France.; The Susanne Levy Gertner Oncogenetics Unit, Chaim Sheba Medical Center, Emek HaEla St 1, Ramat Gan, 52621, Israel.; Sackler Faculty of Medicine, Tel Aviv University, Haim Levanon 30, Ramat Aviv, 69978, Israel.; Department of Surgery, Mount Sinai Hospital, 600 University Avenue, Toronto, ON M5G 1X5, Canada.; Samuel Lunenfeld Research Institute, 600 University Avenue, Toronto, ON M5G 1X5, Canada.; University Health Network Toronto General Hospital, 200 Elizabeth St, Toronto, ON M5G 2C4, Canada.; Schools of Medicine and Public Health, Division of Cancer Prevention & Control Research, Jonsson Comprehensive Cancer Centre, UCLA, 650 Charles Young Drive South, Los Angeles, CA, 90095-6900, USA.; Cancer Risk and Prevention Clinic, Dana-Farber Cancer Institute, 450 Brookline Avenue, Boston, MA, 02215, USA.; Department of Preventive Medicine, Keck School of Medicine, University of Southern California, 1975 Zonal Ave, Los Angeles, CA, 90033, USA.; Center for Cancer Prevention and Translational Genomics, Samuel Oschin Comprehensive Cancer Institute, Cedars-Sinai Medical Center, Spielberg Building, 8725 Alden Dr, Los Angeles, CA, 90048, USA.; Department of Biomedical Sciences, Cedars-Sinai Medical Center, Spielberg Building, 8725 Alden Dr, Los Angeles, CA, 90048, USA.; Cancer Epidemiology & Intelligence Division, Cancer Council Victoria, 615 St Kilda Road, Melbourne, Victoria, 3004, Australia.; Centre for Epidemiology and Biostatistics, Melbourne School of Population and Global Health, The University of Melbourne, Level 1, 723 Swanston Street, Melbourne, Victoria, 3010, Australia.; Department of Epidemiology and Preventive Medicine, Monash University, Melbourne, Victoria,Australia.; Department of Pathology and Laboratory Medicine, University of Kansas Medical Center, 3901 Rainbow Blvd, Kansas City, KS, 66160, USA.; Department of Medicine, McGill University, 1001 Decarie Boulevard, Montréal, QC, H4A3J1, Canada.; Division of Clinical Epidemiology, Royal Victoria Hospital, McGill University, 1001 Decarie Boulevard, Montréal, QC, H4A3J1, Canada.; Department of Dermatology, Huntsman Cancer Institute, University of Utah School of Medicine, 2000 Circle of Hope, Salt Lake City, UT, 84112, USA.; Department of Health Sciences Research, Mayo Clinic, 200 First St. SW, Rochester, MN, 55905, USA.; Cancer Prevention and Control, Samuel Oschin Comprehensive Cancer Institute, Cedars-Sinai Medical Center, 8700 Beverly Blvd., Room 1S37, Los Angeles, CA, 90048, USA.; Community and Population Health Research Institute, Department of Biomedical Sciences, Cedars-Sinai Medical Center, 8700 Beverly Blvd., Room 1S37, Los Angeles, CA, 90048, USA.; Public Health Sciences Division, Swedish Cancer Institute, 1221 Madison St. Ste 300, Seattle, WA, 98109, USA.; Unit of Clinical Chemistry, Department of Medical Biosciences, Umeå University, By 6M van 2, Sjukhusomradet, Umea universitet, Umea, 901 85, Sweden.; Clinical Genetics Branch, DCEG, National Cancer Institute, 9609 Medical Center Dr, Bethesda, MD, 20850-9772, USA.; Cancer & Environment Group, Center for Research in Epidemiology and Population Health (CESP), INSERM, University Paris-Sud, University Paris-Saclay, Villejuif, 94805 France.; Department of Environmental Medicine, Division of Nutritional Epidemiology, Karolinska Institutet, Nobels väg 13, SE-171 77, Stockholm, SE-171, Sweden.; Department of Oncology, Södersjukhuset, Sjukhusbacken 10, 118 83 Stockholm, Sweden.; Molecular Genetics of Breast Cancer, German Cancer Research Center (DKFZ), Im Neuenheimer Feld 580, Heidelberg, 69120, Germany.; Nuffield Department of Surgical Sciences, Faculty of Medical Science, John Radcliffe Hospital, University of Oxford, Oxford OX1 2JD, UK.; Department of Surgical Oncology, Princess Margaret Cancer Centre, 610 University Avenue, Toronto, Ontario, M5G2M9, Canada.; Department of Internal Medicine 1, University Hospital Dresden, Technische Universität Dresden (TU Dresden), 01307 Dresden, Germany.; National Institute of Occupational Health (STAMI), Gydas vei 8, 0033, Oslo, Norway.; Department of Gynecology and Gynecologic Oncology, Dr. Horst Schmidt Kliniken Wiesbaden, Wiesbaden, Germany.; Department of Gynecology and Gynecologic Oncology, Kliniken Essen-Mitte/ Evang. Huyssens-Stiftung/ Knappschaft GmbH, Henricistrasse 92, Essen, 45136, Germany.; Early Detection and Prevention Section, International Agency for Research on Cancer, 150 cours Albert Thomas, Lyon, 69008, France.; Department of Virus, Lifestyle and Genes, Danish Cancer Society Research Center, Strandboulevarden 49, Copenhagen, DK-2100, Denmark.; Molecular Unit, Department of Pathology, Herlev Hospital, University of Copenhagen, Herlev Ringvej 75, Herlev, DK-2730, Denmark.; Preventive Medicine, Seoul National University College of Medicine, 1 Gwanak-ro, Gwanak-gu, Seoul 151 742, Korea.; German Research Center for Environmental Health, Institute for Cancer Research, Ingolstadter Landstr. 1, London, SM2 5NG, UK.; Center for Medical Genetics, NorthShore University HealthSystem, 1000 Central St, Evanston, IL,60201, USA.; The University of Chicago Pritzker School of Medicine, 924 E 57th St, Chicago, IL, 60637, USA.; British Columbia’s Ovarian Cancer Research (OVCARE) Program, Vancouver General Hospital, BC Cancer Agency and University of British Columbia, #3427-600 West 10th Avenue, Vancouver, BC, V5Z 4E6, Canada.; Department of Molecular Oncology, BC Cancer Agency Research Centre, #3427-600 West 10th Avenue, Vancouver, BC, V5Z 4E6, Canada.; Department of Pathology and Laboratory Medicine, University of British Columbia, #3427-600 West 10th Avenue, Vancouver, BC, V5Z 4E6, Canada.; N.N. Petrov Institute of Oncology, Leningradskaya ul., 68, St. Petersburg, 197758, Russia.; Lombardi Comprehensive Cancer Center, Georgetown University, 3800 Reservoir Road, Washington, DC, 20007, USA.; Independent Laboratory of Molecular Biology and Genetic Diagnostics, Pomeranian Medical University, Rybacka 1, 70-204 Szczecin, Poland.; Parkville Familial Cancer Centre, Peter MacCallum Cancer Center, 305 Grattan Street, Melbourne, Victoria, 3000, Australia.; Department of Radiation Sciences, Umeå University, By 6M van 2, Sjukhusomradet, Umea universitet, 901 85, Umea, Sweden.; Department of Medicine, Division of Oncology and Stanford Cancer Institute, Stanford University School of Medicine, 780 Welch Rd, Stanford, CA 94304, USA.; Clinical and Translational Epidemiology Unit, Massachusetts General Hospital, 02114 Boston MA.; Molecular Medicine Center, Department of Medical Chemistry and Biochemistry, Medical Faculty, Medical University of Sofia, Sofia, Bulgaria.; Women’s Cancer Program at the Samuel Oschin Comprehensive Cancer Institute, Cedars-Sinai Medical Center, 8700 Beverly Boulevard, Los Angeles, CA, 90048, USA.; Hollings Cancer Center and Department of Public Health Sciences, Medical University of South Carolina, 68 President Street Bioengineering Building, MSC955, Charleston, SC, 29425, USA.; Cancer Epidemiology, University Cancer Center Hamburg (UCCH), University Medical Center Hamburg-Eppendorf, Martinistraße 52, Hamburg, 20246, Germany.; Clinical Gerontology, Department of Public Health and Primary Care, University of Cambridge, 2 Worts’ Causeway, Cambridge, CB1 8RN, UK.; Department of Genetics and Fundamental Medicine, Bashkir State University, ul. Zaki Validi 32, Ufa, 450076, Russia.; Institute of Biochemistry and Genetics, Ufa Scientific Center of Russian Academy of Sciences, 71 prosp. Oktyabrya, Ufa, 450054, Russia.; Division of Urologic Surgery, Brigham and Womens Hospital, Boston, Massachusettes, 02115, USA.; Radboud Institute for Health Sciences, Radboud University Medical Center, Geert Grooteplein 21, Nijmegen, 6525 EZ, The Netherlands.; Department of Genitourinary Medical Oncology, University of Texas MD Anderson Cancer Center, 1155 Pressler St, Houston, TX, 77030, USA.; Department of Gynaecology, Rigshospitalet, University of Copenhagen, Blegdamsvej 9, Copenhagen, DK-2100, Denmark.; Prosserman Centre for Health Research, Lunenfeld-Tanenbaum Research Institute, Sinai Health System, 60 Murray Street, Toronto, Ontario, M5T 3L9, Canada.; Division of Epidemiology, Dalla Lana School of Public Health, University of Toronto, 155 College Street, Toronto, ON, M5T3M7, Canada.; ISGlobal, Centre for Research in Environmental Epidemiology (CREAL), Barcelona, 08036, Spain.; IMIM (Hospital del Mar Research Institute), Barcelona, Spain.; Universitat Pompeu Fabra (UPF), Barcelona, Spain.; Division of Cancer Epidemiology and Genetics, National Cancer Institute, National Institutes of Health, Department of Health and Human Services, 9609 Medical Center Dr, Bethesda, MD, 20892, USA.; Department of Cancer Genetics, Institute for Cancer Research, Oslo University Hospital Radiumhospitalet, Ullernchausseen 70, Oslo, 0379, Norway.; Institute of Clinical Medicine, Faculty of Medicine, University of Oslo, Kirkeveien 166, Oslo, 0450,Norway.; Department of Clinical Molecular Biology, Oslo University Hospital, University of Oslo, Kirkeveien 166, Oslo, 0450, Norway.; Department of Pathology and Laboratory Diagnostics, the Maria Sklodowska-Curie Institute-Oncology Center, Roentgena 5, Warsaw, 02-781, Poland.; Department of Otorhinolaryngology, Head and Neck Surgery, Maastricht University Medical Center, P. Debyelaan 25, P.O. Box 5800, Maastricht, 6202 AZ, The Netherlands.; Department of Integrative Oncology, British Columbia Cancer Agency, Room 10-111 675 West 10th Avenue, Vancouver, BC, V5Z1L3, Canada.; VIB Center for Cancer Biology, VIB, Herestraat 49, Leuven, 3001, Belgium.; Laboratory for Translational Genetics, Department of Human Genetics, University of Leuven, Oude Markt 13, Leuven, 3000, Belgium.; Integrative Tumor Epidemiology Branch, DCEG, National Cancer Institute, 9609 Medical Center Drive, Room SG/7E106, Rockville, MD 20850, USA.; College of Pharmacy, Washington State University, PBS 431 PO Box 1495 Washington State University, Spokane, WA 99210-1495, USA.; Cancer Control Research, BC Cancer Agency, 675 West 10th Avenue, Vancouver, BC, V5Z 1L3, Canada.; Clalit Health Services, Clalit National Israeli Cancer Control Center, Carmel Medical Center, 2 Horev Street, Haifa, 3436212, Israel.; Institute of Human Genetics, University Medical Center Hamburg-Eppendorf, Martinistraße 52, Hamburg, 20246, Germany.; Gynecology Service, Department of Surgery, Memorial Sloan Kettering Cancer Center, 1275 York Avenue, New York, NY, 10065, USA.; Gynecologic Oncology, Laura and Isaac Pearlmutter Cancer Center, NYU Langone Medical Center,240 East 38th Street 19th Floor, New York, NY, 10016, USA.; Department of Family Medicine and Community Health, Mary Ann Swetland Center for Environmental Health, Case Western Reserve University, Cleveland, OH, 44106, USA.; Servicio Galego de Saude (SERGAS), Instituto de Investigación Sanitaria de Santiago de Compostela (IDIS), Santiago De Compostela, 15706, Spain.; Translational Research Program, Fred Hutchinson Cancer Research Center, Seattle, WA, 98109, USA.; Department of Molecular Medicine and Surgery, Karolinska Institutet, Karolinska Univ Hospital, Stockholm, 171 76, Sweden.; Health Sciences Research, Mayo Clinic Arizona, 13400 E. Shea Blvd, Scottsdale, AZ, 85259, USA.; Epidemiology Division, Princess Margaret Cancer Centre, 610 University Avenue, Toronto, Ontario,M5G2M9, Canada.; Unit of Oncology 1, Department of Clinical and Experimental Oncology, Istituto Oncologico Veneto IRCCS, 35122 Padua Italy.; Department of Medical Genetics, Oslo University Hospital, Kirkeveien 166, Oslo, 0450, Norway.; Institute of Human Genetics, University Hospital Ulm, Prittwitzstrasse 43, Ulm, 89075, Germany.; Translational Cancer Research Area, University of Eastern Finland, Yliopistonranta 1, Kuopio, 70210, Finland.; Institute of Clinical Medicine, Pathology and Forensic Medicine, University of Eastern Finland, Yliopistonranta 1, Kuopio, 70210, Finland.; Imaging Center, Department of Clinical Pathology, Kuopio University Hospital, Puijonlaaksontie 2, Kuopio, 70210, Finland.; Epidemiology Program, University of Hawaii Cancer Center, 701 Ilalo St, Honolulu, HI, 96813, USA.; Department of Clinical Science and Education, Södersjukhuset, Karolinska Institutet, Stockholm, Sweden.; Division of Gynecologic Oncology, University Health Network, Princess Margaret Hospital, 610 University Avenue, OPG Wing, 6-811 Toronto, Ontario, M5G 2M9, Canada.; Division of Gynaecology and Obstetrics, Technische Universität München, Arcisstraße 21, Munich, 80333, Germany.; Faculty of Medicine, University of Heidelberg. In Neuenheimer Feld 672, 69120 Heidelberg, Germany.; NRG Oncology, Statistics and Data Management Center, Roswell Park Cancer Institute, Elm & Carlton Streets, Buffalo, NY, 14263, USA.; Womens Cancer Research Center, Magee-Womens Research Institute and Hillman Cancer Center, Pittsburgh, PA, 15213, USA.; Division of Gynecologic Oncology, Department of Obstetrics, Gynecology and Reproductive Sciences, University of Pittsburgh School of Medicine, 300 Halket Street, Pittsburgh, PA, 15213, USA.; Immunology and Molecular Oncology Unit, Veneto Institute of Oncology IOV-IRCCS, Via Gattamelata 64, Padua, 35128, Italy.; Catalan Institute of Oncology, Bellvitge Biomedical Research Institute (IDIBELL), Consortium for Biomedical Research in Epidemiology and Public Health (CIBERESP) and University of Barcelona, Barcelona, Spain.; Division of Cancer Prevention and Control, Roswell Park Cancer Institute, Elm & Carlton Streets, Buffalo, NY, 14263, USA.; Division of Population Health, Health Services Research and Primary Care, University of Manchester, Oxford Road, Manchester, M13 9PL, UK.; Division of Health Sciences, Warwick Medical School, Warwick University, University of Warwick,Coventry, CV4 7AL, UK.; Department of Laboratory Medicine and Pathobiology, University of Toronto, 1 King’s College Circle, Toronto, ON, M5S1A8, Canada.; Laboratory Medicine Program, University Health Network, 200 Elizabeth Street, Toronto, ON, M5G2C4, Canada.; Department of Medicine, Abramson Cancer Center, Perelman School of Medicine at the University of Pennsylvania, 3400 Civic Center Boulevard, Philadelphia, PA, 19104, USA.; Department of Oncology, Addenbrooke’s Hospital, University of Cambridge, Cambridge, CB1 8RN, UK.; NIHR Bristol Biomedical Research Centre Nutrition Theme, University of Bristol, Upper Maudlin Street, Bristol, BS2 8AE, UK.; Department of Population Sciences, Beckman Research Institute of City of Hope, 1500 E Duarte, CA, 91010, USA.; Department of Obstetrics and Gynecology, Helsinki University Hospital, University of Helsinki, Haartmaninkatu 8, Helsinki, 00290, Finland.; Department of Urology, University of Washington, Seattle, Washington, 98195, USA.; Center for Genomic Medicine, Rigshospitalet, Copenhagen University Hospital, Blegdamsvej 9, Copenhagen, DK-2100, Denmark.; Latvian Biomedical Research and Study Centre, Ratsupites str 1, Riga, Latvia.; Cancer Genetics and Prevention Program, University of California San Francisco, 1600 Divisadero St., San Francisco, CA, 94143-1714, USA.; Clinical Genetics Research Lab, Department of Cancer Biology and Genetics, Memorial Sloan-Kettering Cancer Center, 1275 York Avenue, New York, NY, 10065, USA.; Clinical Genetics Service, Department of Medicine, Memorial Sloan-Kettering Cancer Center, 1275 York Avenue, New York, NY, 10065, USA.; Department of Molecular Genetics, National Institute of Oncology, Ráth György u. 7-9, Budapest, 1122, Hungary.; Department of Clinical Neurosciences, University of Cambridge, Cambridge, CB2 0QQ, UK.; Center for Clinical Cancer Genetics, The University of Chicago, 5841 S Maryland Ave, Chicago, IL, 60637, USA.; Department of Epidemiology, Gillings School of Global Public Health, University of North Carolina, 135 Dauer Dr, Chapel Hill, NC, 27599-7435, USA.; UNC Lineberger Comprehensive Cancer Center, 450 West Dr, Chapell Hill, NC, 27599, USA.; The University of Surrey, Guildford, Surrey, GU2 7XH, UK.; Department of Cancer Epidemiology, H. Lee Moffitt Cancer Center and Research Institute, 12902 Magnolia Drive, Tampa, FL, 33612, USA.; Department of Applied Health Research, University College London, 1-19 Torrington Place, London WC1E 6BT, UK.; Centre for Cancer Genetic Epidemiology, Department of Oncology, Strangeways Laboratory, University of Cambridge, CB1 8RN, UK.; Department of Genetics and Computational Biology, QIMR Berghofer Medical Research Institute, 300 Herston Road, Brisbane, Queensland, 4006, Australia.; Department of Obstetrics and Gynecology, Oregon Health & Science University, 3181 SW Sam Jackson Park Road, L-466, Portland, OR, 97239, USA.; Knight Cancer Institute, Oregon Health & Science University, 3181 SW Sam Jackson Park Road, L-466, Portland, OR, 97239, USA.; Department of Gastroenterology, Radboud University Nijmegen Medical Center, Geert Grooteplein Zuid 10, Internal B.O. Box 433, Nijmegen, 6525 GA, The Netherlands.; Department of Epidemiology, University of Washington School of Public Health, 1959 NE Pacific St, Seattle, WA, 98195, USA.; Research Centre for Genetic Engineering and Biotechnology ‘Georgi D. Efremov’, Macedonian Academy of Sciences and Arts, Boulevard Krste Petkov Misirkov, Skopje, 1000, Republic of Macedonia.; Bristol Dental School, University of Bristol, Lower Maudlin Street, Bristol, BS1 2LY, UK.; Unit of Molecular Bases of Genetic Risk and Genetic Testing, Department of Research, Fondazione IRCCS (Istituto Di Ricovero e Cura a Carattere Scientifico) Istituto Nazionale dei Tumori (INT), Via Giacomo Venezian 1, Milan, 20133, Italy.; Decode genetics, Sturlugata 8, IS-101 Reykjavik, Iceland, Reykjavik, Iceland.; School of Women’s and Children’s Health, Faculty of Medicine, University of NSW Sydney, 18 High St, Sydney, New South Wales, 2052, Australia.; The Kinghorn Cancer Centre, Garvan Institute of Medical Research, 384 Victoria Street, Sydney, New South Wales, 2010, Australia.; Division of Epidemiology, Department of Medicine, Vanderbilt Epidemiology Center, Vanderbilt-Ingram Cancer Center, Vanderbilt University School of Medicine, 1161 21st Ave S # D3300, Nashville, TN, 37232, USA.; Clalit National Cancer Control Center, Carmel Medical Center and Technion Faculty of Medicine, 7 Michal Street, Haifa 34362, Israel.; Department of Genetics, University of Pretoria, Private Bag X323, Arcadia, 0007, South Africa.; Department of Chronic Disease Epidemiology, Yale School of Public Health, 60 College St, New Haven, CT, 06510, USA.; Cancer Center Cluster Salzburg at PLUS, Department of Molecular Biology, University of Salzburg, Billrothstr.11, 5020 Salzburg, Austria.; Division of Epigenomics and Cancer Risk Factors, DKFZ – German Cancer Research Center, Im Neuenheimer Feld 280, 69120 Heidelberg, Germany.; Translational Lung Research Center Heidelberg (TLRC-H), Member of the German Center for Lung Research (DZL), Heidelberg, 69120, Germany.; Department of Urology, Erasmus University Medical Center, Wytemaweg 80, Rotterdam, 3015 CN, The Netherlands.; Department of Radiation Oncology, Icahn School of Medicine at Mount Sinai, 1425 Madison Avenue, New York, NY, 10029, USA.; Department of Genetics and Genomic Sciences, Icahn School of Medicine at Mount Sinai, 1425 Madison Avenue, New York, NY, 10029, USA.; Department of Epidemiology, University of Washington, M4 C308, 1100 Fairview Ave N, Seattle, WA, 98109, USA.; Faculty of Medicine and Health Sciences, Basic Medical Sciences, Ghent University, De Pintelaan 185, Gent, 9000, Belgium.; Hereditary Cancer Clinic, University Hospital of Heraklion, Voutes, Heraklion, 711 10, Greece.; Epidemiology Branch, National Institute of Environmental Health Sciences, NIH, 111 T.W. Alexander Drive, Research Triangle Park, NC, 27709, USA.; Research Oncology, Guy’s Hospital, King’s College London, Guy’s Hospital Great Maze Pond, London, SE1 9RT, UK.; Institute of Biomedicine, University of Turku, Turku, 20014, Finland.; Division of Laboratory, Department of Medical Genetics, Turku University Hospital, Turku, 20014, Finland.; Prostate Cancer Research Center, Faculty of Medicine and Life Sciences and BioMediTech Institute, University of Tampere, Tampere, 33014, Finland.; Division of Molecular Pathology, The Netherlands Cancer Institute-Antoni van Leeuwenhoek Hospital, Plesmanlaan 121, Amsterdam, 1066 CX, The Netherlands.; Division of Psychosocial Research and Epidemiology, The Netherlands Cancer Institute-Antoni van Leeuwenhoek hospital, Plesmanlaan 121, Amsterdam, 1066 CX, The Netherlands.; Department of Preventive Medicine, Keck School of Medicine, University of Southern California, 1450 Biggy Street, Los Angeles, CA, 90033, USA.; Department of Epidemiology and Biostatistics, Jiangsu Key Lab of Cancer Biomarkers, Prevention and Treatment, Collaborative Innovation Center for Cancer Personalized Medicine, School of Public Health, Nanjing Medical University, 101 Longmian Ave, Jiangning District, Nanjing, 211166, Peoples Republic of China.; Department of Genetics and Genomic Sciences, Department of Population Health Science and Policy, Icahn School of Medicine at Mount Sinai, 1425 Madison Avenue, 2nd floor, New York, NY, 10029, USA.; Dept of OB/GYN and Comprehensive Cancer Center, Medical University of Vienna, Waehringer Guertel 18-20, Vienna, 1090, Austria.; Department of Internal Medicine, University of Utah Health Sciences Center, 295 Chipeta Way,Salt Lake City, UT 84132, USA.; Department of Molecular Medicine, Aarhus University Hospital, Aarhus, DK-8200, Denmark.; Department of Clinical Medicine, Aarhus University, Aarhus, DK-8200, Denmark.; Precision Medicine, School of Clinical Sciences at Monash Health, Monash University, 246 Clayton Road, Clayton, Victoria, 3168, Australia.; Department of Clinical Pathology, The University of Melbourne, Cnr Grattan Street and Royal Parade, Melbourne, Victoria, 3010, Australia.; Department of Medicine III, University Hospital, LMU Munich, Marchioninistr. 15, 81377 Munich, Germany.; The Curtin UWA Centre for Genetic Origins of Health and Disease, Curtin University and University of Western Australia, 35 Stirling Hwy, Perth, Western Australia, 6000, Australia.; Department of Obstetrics and Gynecology, Sahlgrenska Cancer Center, Inst Clinical Scienses, University of Gothenburg, Blå stråket 6, Gothenburg, 41345, Sweden.; Epidemiology Center, College of Medicine, University of South Florida, 3650 Spectrum Blvd., Suite 100, Tampa, FL, 33612, USA.; Division of Breast Cancer Research, The Institute of Cancer Research, London, SW7 3RP, UK.; Department of Molecular Biology, School of Medicine of São José do Rio Preto, Av Brig Faria Lima 5416 Vila São Pedro, São José do Rio Preto, SP, 15090-000, Brazil.; Department of Genetics and Evolutive Biology, Institute of Biosciences, University of São Paulo, Rua do Matão, 321, São Paulo, SP, 05508-090, Brazil.; SWOG Statistical Center, Fred Hutchinson Cancer Research Center, Seattle, Washington, 98109, USA.; Faculty of Medicine, University of Oviedo and CIBERESP, Campus del Cristo s/n, 33006, Oviedo, Spain.; Epigenetic and Stem Cell Biology Laboratory, National Institute of Environmental Health Sciences, NIH, 111 T.W. Alexander Drive, Research Triangle Park, NC, 27709, USA.; Medical Statistics Group, School of Health and Related Research (ScHARR), University of Sheffield, Regent Court, 30 Regent Street, Sheffield, S1 4DA, UK.; Department of Genetics, Portuguese Oncology Institute, Rua Dr. António Bernardino de Almeida 62, Porto, 4220-072, Portugal.; Biomedical Sciences Institute (ICBAS), University of Porto, R. Jorge de Viterbo Ferreira 228, Porto, 4050-013, Portugal.; Department of Epidemiology, Mailman School of Public Health, Columbia University, 722 West 168th Street, New York, NY, 10032, USA.; Obstetrics and Gynecology Epidemiology Center, Brigham and Women’s Hospital, 221 Longwood Avenue RFB 368, Boston, MA, 02115, USA.; Harvard T.H. Chan School of Public Health, 221 Longwood Avenue RFB 368, Boston, MA, 02115, USA.; Department of Clinical Genetics, Odense University Hospital, Sonder Boulevard 29, Odence C, 5000, Denmark.; Department of Gynecology and Obstetrics, Haukeland University Hospital, Bergen, 5021, Norway.; Centre for Cancer Biomarkers CCBIO, Department of Clinical Science, University of Bergen, Bergen, 5021, Norway.; Program in Cancer Genetics, Departments of Human Genetics and Oncology, McGill University, 1001 Decarie Boulevard, Montréal, QC, H4A3J1, Canada.; Department of Medical Genetics, Cambridge University, Hills Road, Cambridge, CB2 0QQ, UK.; Department of Cancer Biology and Genetics, The Ohio State University, 460 W. 12th Avenue, Columbus, OH, 43210, USA.; Institute of Human Genetics, Pontificia Universidad Javeriana, Carrera 7 No. 40-90, Bogota, Colombia.; Division of Cancer Sciences, Manchester Cancer Research Centre, Faculty of Biology, Medicine and Health, Manchester Academic Health Science Centre, NIHR Manchester Biomedical Research Centre, Health Innovation Manchester, University of Manchester, Manchester, M20 4GJ, UK.; Cancer Epidemiology Unit, Nuffield Department of Population Health, University of Oxford, Oxford, OX3 7LF, UK.; Department of Medical Oncology, Beth Israel Deaconess Medical Center, 330 Brookline Avenue, Boston, MA, 02215, USA.; Huntsman Cancer Institute and Department of Population Health Sciences, University of Utah, 2000 Circle of Hope, Rm 4125, Salt Lake City, UT, 84112, USA.; Department of Oncology, Cross Cancer Institute, University of Alberta, 116 St & 85 Ave, Edmonton, AB T6G 2R3, Canada.; Division of Radiation Oncology, Cross Cancer Institute, University of Alberta, 116 St & 85 Ave, Edmonton, AB T6G 2R3, Canada.; Division of Gynecologic Oncology, Department of Obstetrics and Gynaecology and Leuven Cancer Institute, University Hospitals Leuven, Herestraat 49, Leuven, 3000, Belgium.; Fundación Pública Galega Medicina Xenómica & Instituto de Investigación Sanitaria de Santiago de Compostela, calle Choupana s/n, Santiago De Compostela, 15706, Spain.; Population Health Department, QIMR Berghofer Medical Research Institute, 300 Herston Road, Brisbane, Queensland, 4006, Australia.; Biostatistics and Computational Biology Branch, National Institute of Environmental Health Sciences, NIH, 111 T.W. Alexander Drive, Research Triangle Park, NC, 27709, USA.; Department of Otolaryngology/Head and Neck Surgery, University of North Carolina at Chapel Hill, Chapel Hill, NC, USA.; City of Hope Clinical Cancer Genomics Community Research Network, 1500 East Duarte Road, Duarte, CA, 91010, USA.; Division of Cancer Sciences, University of Manchester, Manchester Cancer Research Centre, Manchester Academic Health Science Centre, The Christie Hospital NHS Foundation Trust, Manchester, M13 9PL, UK.; Fred Hutchinson Cancer Research Center, 1100 Fairview Ave N, Seattle, WA, 98109, USA.; Department of Epidemiology, University of Washington, 1100 Fairview Ave N, Seattle, WA, 98109, USA.; Department of Health Research and Policy-Epidemiology, Stanford University School of Medicine, 259 Campus Drive, Stanford, CA, 94305, USA.; Department of Biomedical Data Science, Stanford University School of Medicine, 259 Campus Drive, Stanford, CA, 94305, USA.; Institute of Medical Informatics, Biometry and Epidemiology, Chair of Epidemiology, Ludwig Maximilians University, Neuherberg D-85764, Munich, Bavaria, Germany.; Helmholtz Zentrum Munchen, German Research Center for Environmental Health (GmbH), Institute of Epidemiology, Ingolstadter Landstr. 1, 85764 Neuherberg, Germany.; Institute of Medical Statistics and Epidemiology, Technical University Munich, Germany, Munich,Germany.; Laboratory of Cancer Genetics and Tumor Biology, Cancer and Translational Medicine Research Unit, Biocenter Oulu, University of Oulu, Aapistie 5A, 90220 Oulu, Finland.; Laboratory of Cancer Genetics and Tumor Biology, Northern Finland Laboratory Centre Oulu, Aapistie 5A, 90220 Oulu, Finland.; Department of Surgical Sciences, Uppsala University, 751 85 Uppsala, Sweden.; Academic Unit of Clinical Oncology, University of Sheffield, Weston Park Hospital, Whitham Road, Sheffield, S10 2SJ, UK.; Discipline of Genetics, Memorial University of Newfoundland, St. John’s, NL, A1C 5S7, Canada.; Department of Epidemiology, Division of Cancer Prevention and Population Science, The University of Texas MD Anderson Cancer Center, 1515 Holcombe Blvd, Houston, TX, 77030, USA.; Magee-Womens Hospital, University of Pittsburgh School of Medicine, 300 Halket St, Pittsburgh,PA, 15213, USA.; Center for Genomic Medicine and Department of Anasthesia, Massachusetts General Hospital, Boston, MA 02114, USA.; Human Genetics, Graduate School of Public Health, University of Pittsburgh, UPMC Cancer Pavilion, Suite 4C, Office # 467, 5150 Centre Avenue, Pittsburgh, PA, 15232, USA.; UPMC Hillman Cancer Center, Pittsburgh, PA, USA.; Genetic Cancer Susceptibility Group, International Agency for Research on Cancer, 150 cours Albert Thomas, Lyon, 69008, France.; Oncogenetics Team, The Institute of Cancer Research and Royal Marsden NHS Foundation Trust, Downs Road, Sutton, SM2 5NG, UK.; Genomics Center, Centre Hospitalier Universitaire de Québec-Université Laval Research Center, 2705 Laurier Boulevard, Québec City, QC, G1V4G2, Canada.; UCLA Path and Lab Med, University of California, 10833 Le Conte Ave, Los Angeles, CA, 190095,USA.; Department of Medicine, Epidemiology Section, Institute for Clinical and Translational Research, Baylor Medical College, One Baylor Plaza, MS: BCM451, Suite 100D, Houston, TX 77030-3411, USA.

## Abstract

Quantifying the genetic correlation between cancers can provide important insights into the mechanisms driving cancer etiology. Using genome-wide association study summary statistics across six cancer types based on a total of 296,215 cases and 301,319 controls of European ancestry, we estimate the pair-wise genetic correlations between breast, colorectal, head/neck, lung, ovary and prostate cancer, and between cancers and 38 other diseases. We observed statistically significant genetic correlations between lung and head/neck cancer (*r*_*g*_=0.57, p=4.6×10^−8^), breast and ovarian cancer (*r*_*g*_=0.24, p=7×10^−5^), breast and lung cancer (*r*_*g*_=0.18, p=1.5×10^−6^) and breast and colorectal cancer (*r*_*g*_=0.15, p=1.1×10^−4^). We also found that multiple cancers are genetically correlated with non-cancer traits including smoking, psychiatric diseases and metabolic characteristics. Functional enrichment analysis revealed a significant excess contribution of conserved and regulatory regions to cancer heritability. Our comprehensive analysis of cross-cancer heritability suggests that solid tumors arising across tissues share in part a common germline genetic basis.

## INTRODUCTION

Inherited genetic variation plays an important role in cancer etiology. Large twin studies have demonstrated an excess familial risk for cancer sites including, but not limited to, breast, colorectal, head/neck, lung, ovary and prostate with heritability estimates ranging between 9% (head/neck) to 57% (prostate).^1-3^ Data from nation-wide and multi-generation registries further show that elevated cancer risks go beyond nuclear families and isolated types, as family history of a specific cancer can increase risk for other cancers.^4-6^ Additional evidence for a shared genetic component have been demonstrated by cross-cancer genome-wide association study (GWAS) meta-analyses, which set out to identify genetic variants associated with more than one cancer type. Fehringer *et al.* studied breast, colorectal, lung, ovarian and prostate cancer, and identified a novel locus at 1q22 associated with both breast and lung cancer.^7^ Kar *et al.* focused on three hormone-related cancers (breast, ovarian and prostate), and identified seven novel susceptibility loci shared by at least two cancers.^8^

Previous attempts to estimate the genetic correlation across cancers using GWAS data^9-12^ have mostly relied on restricted maximum likelihood (REML) implemented in GCTA (genome-wide complex trait analysis)^13^ and individual-level genotype data. However, these studies have had limited sample sizes, yielding inconclusive results. Sampson *et al.* quantified genetic correlations across 13 cancers in European ancestry populations and identified four cancer pairs with nominally significant genetic correlations (bladder-lung, testis-kidney, lymphoma-osteosarcoma, lymphoma-leukemia).^9^ They did not observe any significant genetic correlations across common solid tumors including cancers of the breast, lung and prostate.^9^ REML becomes computationally challenging for large sample sizes and is sensitive to technical artifacts. LD score regression (LDSC)^14,15^ overcomes these issues by leveraging the relationship between association statistics and LD patterns across the genome. We recently used cross-trait LDSC to quantify genetic correlations across six cancers based on a subset of the data included here and found moderate correlations between colorectal and pancreatic cancer as well as between lung and colorectal cancer.^16^ However, the average sample size was only 11,210 cases and 13,961 controls per cancer, resulting in imprecise estimates with wide confidence intervals.

In addition to the development of novel analytical methods tailored to genomic data, several high-quality functional annotations have recently been released into the public domain through large-scale efforts. For example, the ENCODE consortium has built a comprehensive and informative parts list of functional elements in the human genome (http://www.nature.com/encode/#/threads), which allows for the analysis of components of SNP-heritability to unravel the functional architecture of complex traits.

Here, we use summary statistics from the largest-to-date European ancestry GWAS of breast, colorectal, head/neck, lung, ovary and prostate cancer with an average sample size of 49,369 cases and 50,219 controls per cancer, to quantify genetic correlations between cancers and their subtypes. We also use GWAS summary statistics for 38 non-cancer traits (average N=113,808 per trait), to quantify the genetic correlations between the six cancers and other diseases. Furthermore, we assessed the proportion of cancer heritability attributable to specific functional categories, with the goal of identifying functional elements that are enriched for SNP-heritability.

Our comprehensive analysis identifies statistically significant genetic correlations between lung and head/neck cancer, breast and ovarian cancer, breast and lung cancer and breast and colorectal cancer. We also find multiple cancers to be genetically correlated with non-cancer traits including smoking, psychiatric diseases and metabolic traits. Functional enrichment analysis reveals a significant contribution of conserved and regulatory regions to cancer heritability. Our results suggest that solid tumors arising across tissues share in part a common germline genetic basis.

## RESULTS

### Heritability estimates across cancers

We first estimated cancer-specific heritability causally explained by common SNPs 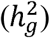 using LDSC (note that this quantity is slightly different from the 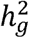 as defined in Yang *et al.*^17^ which estimates the heritability due to genotyped and imputed SNPs) (**see Methods**). Estimates of 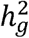 on the liability scale ranged from 0.03 (ovarian) to 0.25 (prostate) (Supplementary Table 1). After removing genome-wide significant (p<5×10^−8^) loci, defined as all SNPs within 500kb of the most significant SNP in a given region (Supplementary Table 2), we observed an ~50% decrease in SNP-heritability for prostate and breast cancer, and ~20% decrease for lung, ovarian and colorectal cancer, despite the fact that we were only excluding 1% (colorectal cancer) to 5% (breast cancer) of the genome. In contrast, the SNP-heritability for head/neck cancer was not affected by removing genome-wide significant loci (Fig. 1A). For most of the cancers, the GWAS significant loci for that particular cancer explained most of the heritability. For some cancers, however, significant GWAS loci of other cancers also explained a non-trivial part of its heritability. For example, the significant breast cancer GWAS loci explained 10%, 15% and 22% heritability of colorectal, ovarian and prostate cancer, respectively; the significant colorectal cancer GWAS loci explained 11% heritability of prostate cancer; the significant lung cancer GWAS loci explained 10% heritability of head/neck cancer; and the significant prostate cancer GWAS loci explained 11% and 15% heritability of breast and ovarian cancer, respectively (Supplementary Table 3). Comparing the liability-scale SNP-heritability to corresponding estimates from twin studies suggests that common SNPs can almost entirely explain the classical heritability of head/neck cancer, whereas for other cancers, only 30–40% of heritability can be explained (Fig. 1B).

**Figure 1.**
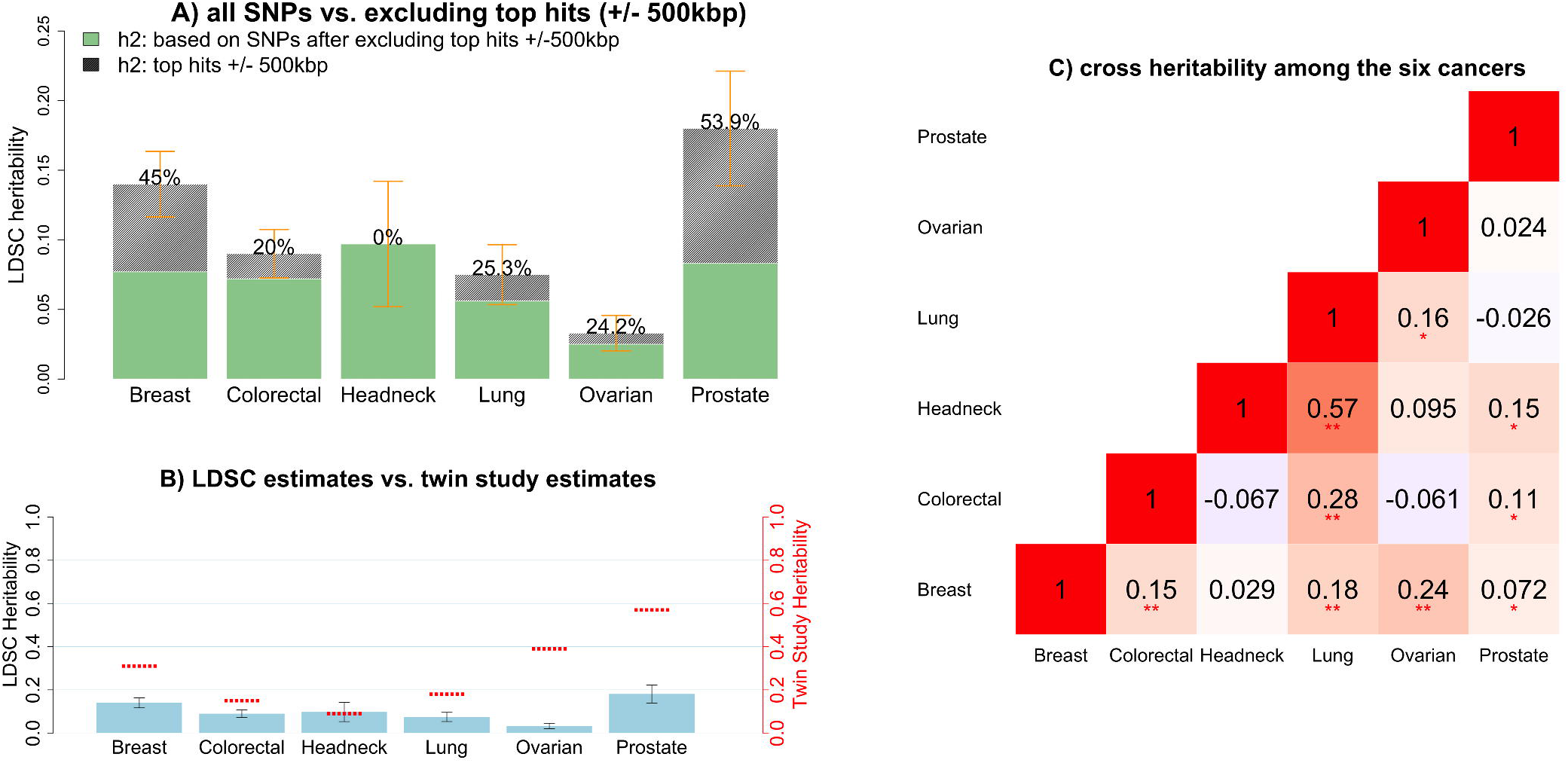
Estimates of SNP-heritability 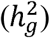 and cross-cancer heritability (*r*_*g*_) based on HapMap3 SNPs calculated using LD score regression (LDSC) for the six cancer types. **A):** the solid bar represents overall SNP 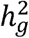 on the liability scale, calculated based on all HapMap3 SNPs. The dark green bar represents 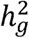 calculated based on “non-significant” SNPs-the remaining SNPs after excluding genome-wide significant hits (p<5×10^−8^) ± 500kb. The black bar with density texture indicates proportion of**/ij** (as reflected by the percentages displayed on top of each bar) that could be explained by top hits ± 500kb surrounded areas. The orange error bars represent 95% confidence intervals. **B):** the solid blue bar represents overall SNP 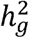 in liability scale (no SNP exclusion), with black error bars indicating 95% confidence intervals. The red short lines correspond to classical estimates of *h*^2^ measured in a twin study of Scandinavian countries (Mucci et al. *JAMA.* 2016;315(1):68). **C):** genetic correlations between cancers. Estimates withstood Bonferroni corrections (p<0.05/15) are marked with double stars (**), and nominal significant results (p<0.05) are marked with single star (*).

### Genetic correlations between cancers

We then estimated the genetic correlation between cancers using cross-trait LDSC (**see Methods**). After adjusting for the number of tests (p<0.05/15=0.003), we found multiple significant genetic correlations Fig. 1C, Supplementary Table 1), with the strongest result observed for lung and head/neck cancer (*r*_*g*_=0.57, se=0.10). In addition, colorectal and lung cancer (*r*_*g*_=0.28, se=0.06), breast and ovarian cancer (*r*_*g*_=0.24, se=0.06), breast and lung cancer (*r*_*g*_=0.18, se=0.04), and breast and colorectal cancer (*r*_*g*_=0.15, se=0.04) showed statistically significant genetic correlations. We also observed nominally significant genetic correlations (p<0.05) between lung and ovarian cancer (*r*_*g*_=0.16, se=0.08), prostate cancer and head/neck (*r*_*g*_=0.15, se=0.08), colorectal (*r*_*g*_=0.11, se=0.05) and breast cancer (*r*_*g*_=0.07, se=0.03) (Fig. 1C). Some cancer pairs showed minimal correlations with estimates close to 0 (ovarian and prostate: *r*_*g*_=0.02, se=0.07; lung and prostate: ***r_g_**=* −0.03, se=0.04; breast and head/neck: *r*_*g*_=0.03, se=0.06). We further calculated the cross-cancer genetic correlation based on data after excluding the GWAS significant regions of each cancer. The estimates were mostly consistent with the results calculated based on all SNPs (data not shown).

We conducted subtype-specific analysis for breast, lung, ovarian and prostate cancer (Supplementary Table 1). Estrogen receptor (ER)+ and ER-breast cancer showed a genetic correlation of 0.60 (se=0.03), indicating that the genetic contributions to these two subtypes are in part distinct. The genetic correlation between the two common lung cancer subtypes adenocarcinoma and squamous cell carcinoma was similarly 0.58 (se=0.10). Further, we observed a significantly larger genetic correlation of lung cancer with ER-(*r*_*g*_=0.29, se=0.06) than with ER+ breast cancer (*r*_*g*_=0.13, se=0.04) (p_difference_=0.002). This also held true for lung squamous cell carcinoma, which showed statistically stronger genetic correlation with ER-(*r*_*g*_=0.33, se=0.08) than with ER+ breast cancer (*r*_*g*_=0.11, se=0.05) (p_difference_=0.0019). We observed no other statistically significant differential genetic correlations across subtypes (all p_difference_>0.1).

We then estimated *local* genetic correlations between cancers using p-HESS, dividing the genome into 1,703 regions (**see Methods**) (Fig. 2 and Supplementary Fig 1). We found that although the genome-wide genetic correlation between breast and prostate cancer was modest (*r*_*g*_=0.07), chr10:123M (10q26.13, p=1.0×10^−7^) and chr9:20-22M (9p21, p=1.0×10^−6^), two previously known pleiotropic regions^18^, showed significant genetic correlations (*r*_*g*_= −0.00098 and *r*_*g*_= 0.00046). Similarly, although the genome-wide genetic correlation between lung and prostate cancer was negligible (*r*_*g*_= −0.03), two previously identified pleiotropic regions (chr6:30-31M or 6p21.33, p=5.7×10^−7^ and chr20:62M or 20q13.33, p=2.8×10^−6^) exhibited significant local genetic correlations (*r*_*g*_= −0.00060 and *r*_*g*_= 0.00067). Overall, local genetic correlation analysis reinforced shared effects for 44% (31/71) of previously reported pleiotropic cancer regions (Supplementary Table 4). It also identified novel pleiotropic signals. For example, the breast and prostate cancer pleiotropic region at 2q33.1 showed significant local genetic correlation between breast and ovarian cancer (p=2.3×10^−6^). Additionally, 6p21.32, a region indicated for head/neck and prostate cancer, showed highly significant local genetic correlation for head/neck and lung cancer (p=8.6×10^−8^).

**Figure 2.**
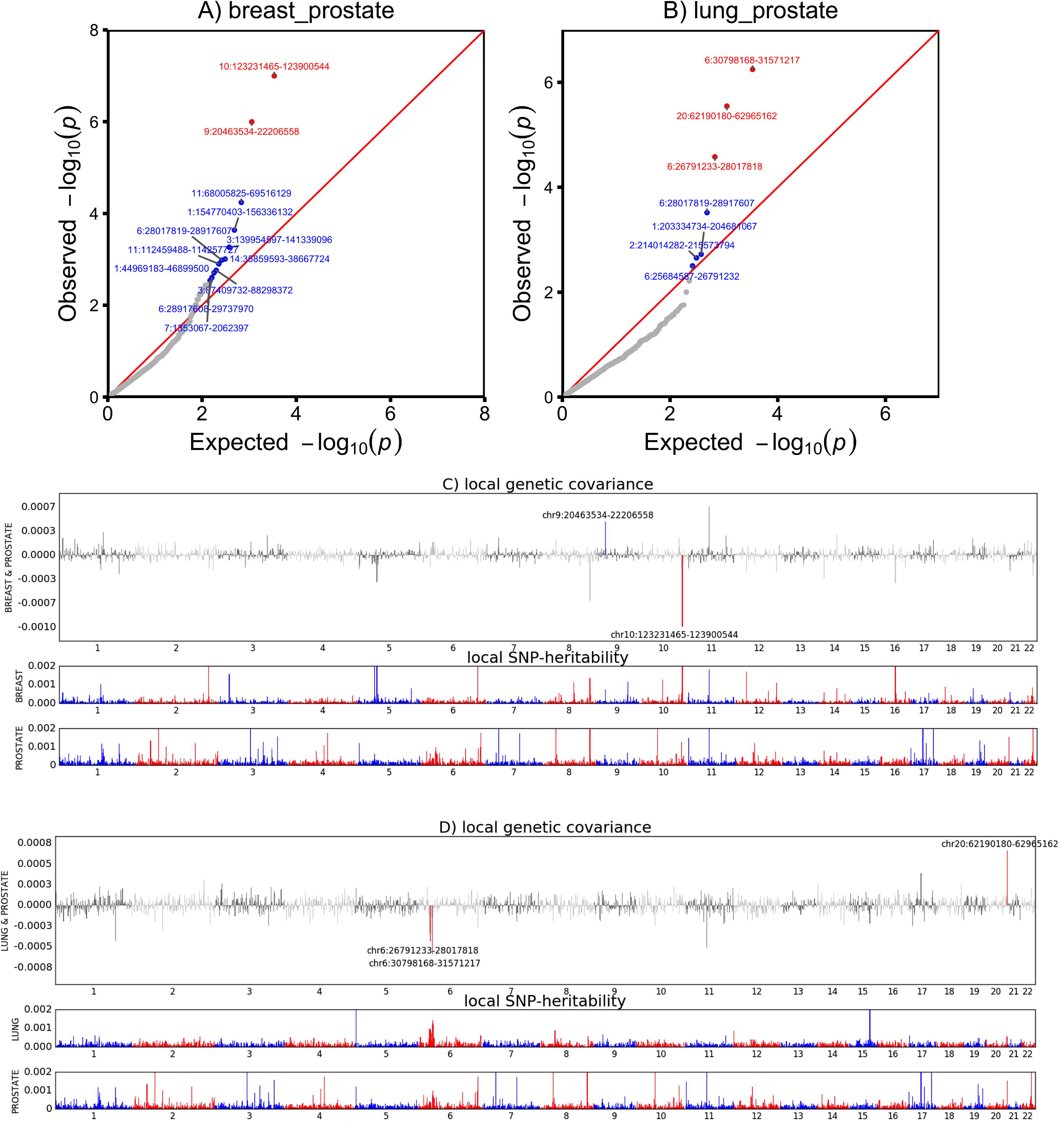
QQ-plots showing region-specific p-values for the local genetic covariance for breast and prostate cancer **(A)**, and for lung and prostate cancer **(B)**. Each dot presents a specific genomic region. In the QQ plots, red color indicates significance after multiple corrections (p<0.05/1,703 regions compared), and blue color indicates nominal significance (p<0.05/15 pairs of cancers compared). Manhattan-style plots showing the estimates of local genetic covariance for breast and prostate cancer **(C)**, and for lung and prostate cancer **(D).** Although breast and prostate cancer only show modest genome-wide genetic correlation, two loci exhibit significant local genetic covariance. Similarly, albeit the negligible overall genetic correlation for lung and prostate cancer, three loci present significant local genetic covariance. In the Manhattan plots, red color indicates even number chromosomes and blue color indicates odd number chromosomes.

### Genetic correlations between cancer and other traits

Significant genetic correlations (p<0.05/228=0.0002) between the six cancers and 38 non-cancer traits reflected several known associations (Fig. 3 and Supplementary Table 5). We observed a strong genetic correlation between smoking and lung cancer (*r*_*g*_=0.56, se=0.06), and similarly for head/neck cancer (*r*_*g*_=0.47, se=0.08), both cancers having smoking as its primary risk factor. ^9^, Educational attainment was negatively genetically correlated with colorectal (*r*_*g*_= −0.17, se=0.04), head/neck (*r*_*g*_= −0.42, se=0.07) and lung cancer (*r*_*g*_= −0.39, se=0.04) (all p<5×10^−6^). Body mass index (BMI) showed a positive genetic correlation with colorectal cancer (*r*_*g*_=0.15, se=0.03) and also suggestive but weak negative correlations with prostate (*r*_*g*_= −0.07, se=0.03) and breast cancer (*r*_*g*_= −0.06, se=0.03). Lung cancer showed a negative genetic correlation with lung function (*r*_*g*_ = −0.15, se=0.04) and age at natural menopause (*r*_*g*_= −0.25, se=0.05), and moderate positive genetic correlations with depressive symptoms (*r*_*g*_=0.25, se=0.06) and waist-to-hip ratio (*r*_*g*_=0.16, se=0.04). Breast cancer showed a positive genetic correlation with schizophrenia (*r*_*g*_=0.14, se=0.03).

**Figure 3.**
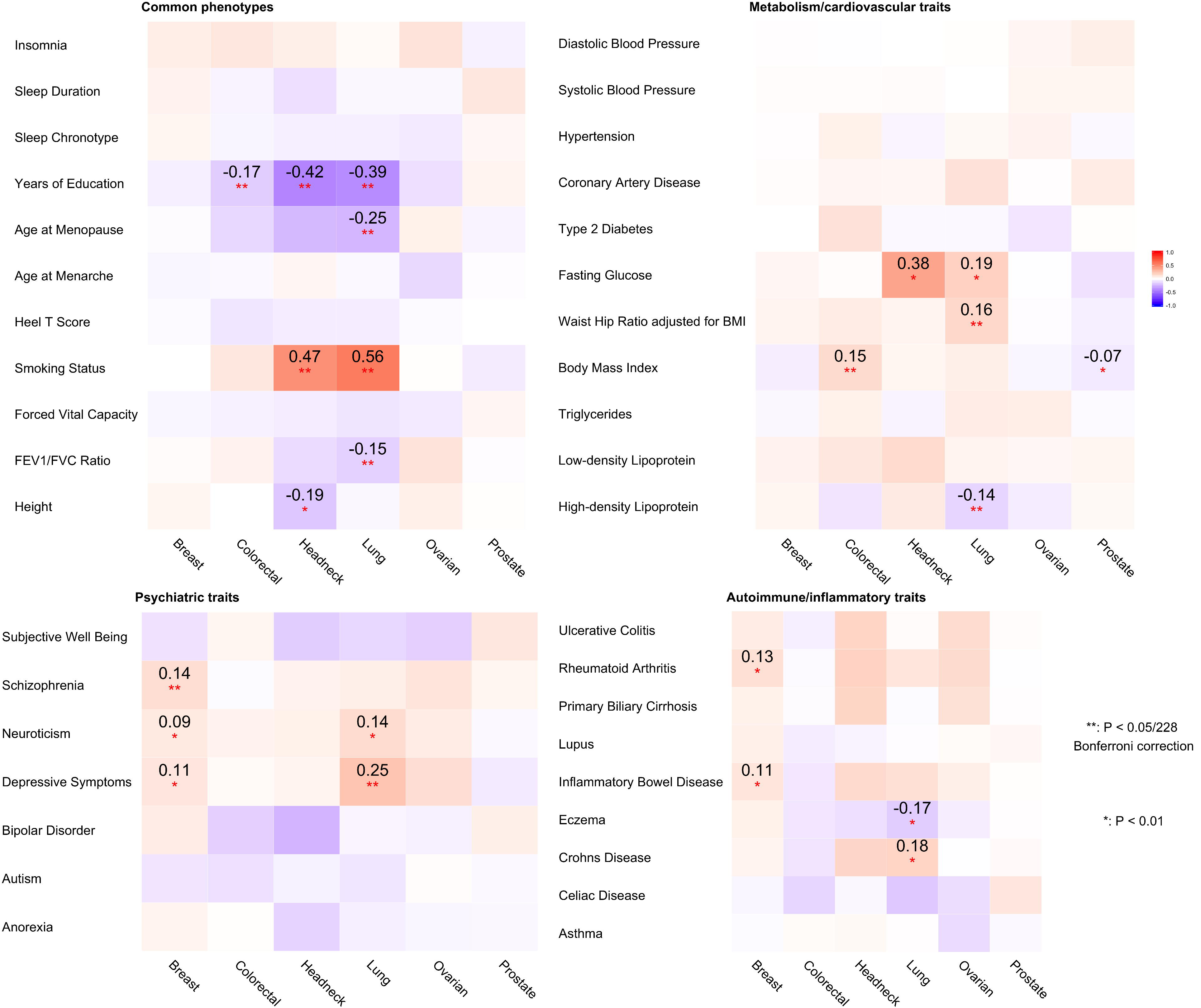
Cross-trait genetic correlation (*r*_*g*_) analysis between six cancers and thirty-eight non-cancer traits. The traits were divided into four categories: **A)** Common phenotypes, **B)** Metabolic or cardiovascular related traits, **C)** Psychiatric traits, **D)** Autoimmune inflammatory diseases. Pairwise genetic correlations withstood Bonferroni corrections (228 tests) are marked with double stars (**), with estimates of correlation shown in the cells. Pairwise genetic correlations with significance at p<0.01 are marked with a single star (*). The color of cells represents the magnitude of correlation.

We did not find evidence of genetic correlations between cancer and several previously suggested risk factors^21-23^ including cardiovascular traits (coronary artery disease, hypertension and blood pressure) or sleep characteristics (chronotype, duration and insomnia). Further, we did not observe genetic correlations between cancer and circulating lipids (HDL, LDL, triglycerides) or type 2 diabetes-related traits except a significant negative correlation between HDL and lung cancer (*r*_*g*_= −0.14, se=0.04). We observed no significant genetic correlation between breast cancer and age at menarche (*r*_*g*_ = −0.03, se=0.03) or age at natural menopause (*r*_*g*_= −0.01, se=0.03). We also did not observe notable genetic correlations between cancer and autoimmune inflammatory diseases or height.

Subtype analysis revealed that smoking and educational attainment showed genetic correlations with all lung cancer subtypes (Supplementary Table 5). Educational attainment, forced vital capacity and depressive symptoms showed genetic correlations with ER-but not ER+ breast cancer, whilst the observed genetic correlation between schizophrenia and breast cancer was limited to ER+ disease, and the genetic correlation between depressive symptoms and lung cancer was observed only for lung squamous cell carcinoma.

We further assessed the support for mediated or pleiotropic causal models for non-cancer traits and cancer using the correlation between trait-specific effect sizes of genome-wide significant SNPs for pairs of phenotypes. We detected four putative directional genetic correlations (defined as p<0.05 from a likelihood ratio (LR) comparing the best non-causal model to the best causal model) (Fig. 4), where SNPs associated with the non-cancer trait showed correlated effect estimates with cancer but the reverse was not true (circulating HDL concentrations and breast cancer, LR_non-causal vs. causal_=0.04, schizophrenia and breast cancer, LR_non-causal vs. causal_=0.003, age at natural menopause and breast cancer, LR_non-causal vs. causal_=0.04, lupus and prostate cancer, LR_non-causal vs. causal_=0.0006).

**Figure 4.**
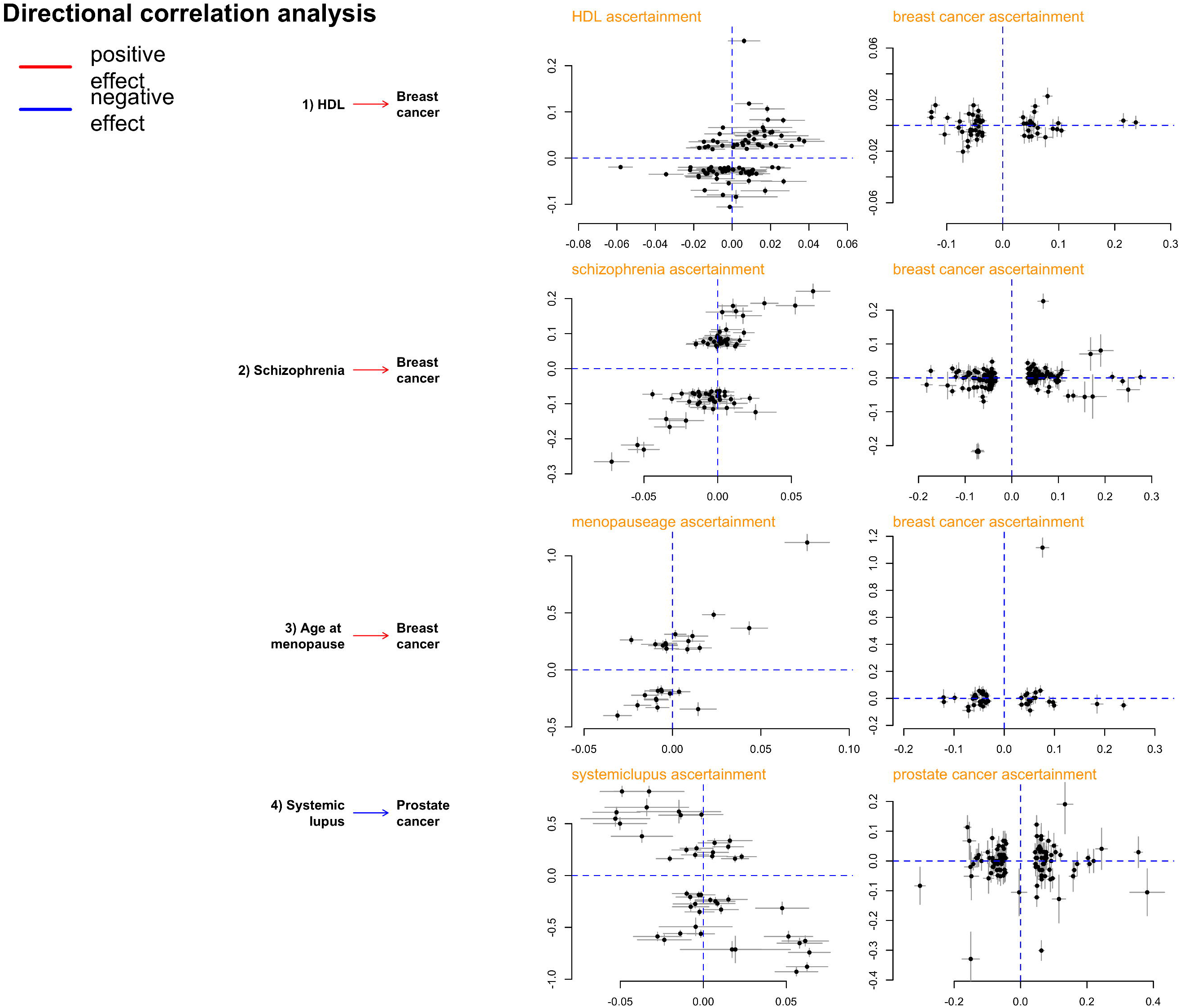
Putative directional relationships between cancers and traits. For each cancer-trait pair identified as candidates to be related in a causal manner, the plots show trait-specific effect sizes (beta coefficients) of the included genetic variants. Grey lines represent the relevant standard errors. **A)** HDL and breast cancer. Trait-specific effect sizes for HDL and breast cancer are shown for SNPs associated with HDL levels (left) and breast cancer (right). **B)** Schizophrenia and breast cancer. Trait-specific effect sizes for HDL and breast cancer are shown for SNPs associated with schizophrenia (left) and breast cancer (right). **C)** Age at natural menopause and breast cancer. Trait-specific effect sizes for age at natural menopause and breast cancer are shown for SNPs associated with age at natural menopause (left) and breast cancer (right). **D)** Lupus and prostate cancer. Trait-specific effect sizes for lupus and prostate cancer are shown for SNPs associated with lupus (left) and prostate cancer (right).

### Functional enrichment analysis of cancer heritability

Finally, we partitioned SNP-heritability of each cancer by using 24 genomic functional annotations (the baseline-LD model described in Gazal *et al.*^24^) and 220 cell-type-specific histone mark annotations (the cell-type-specific model described in Finucane *et al.*^14^). Meta-analysis across the six cancers revealed statistical significant enrichments for multiple functional categories. We observed the highest enrichment for conserved regions (Table 1, Supplementary Table 6) which overlapped with only 2.6% of SNPs but explained 25% of cancer SNP-heritability (9.8-fold enrichment, p=2.3×10^−5^). Transcription factor binding sites showed the second highest enrichment (4.0-fold, 13% of SNPs explaining 40% of SNP-heritability, p=1.4×10^−7^). Further, super-enhancers (groups of putative enhancers in close genomic proximity with unusually high levels of mediator binding) showed a significant 2.6-fold enrichment (p=2.0×10^−20^). Additional enhancers, including regular enhancers (3.2-fold), weak enhancers (3.1-fold) and FANTOM5 enhancers (3.1-fold), presented similar enrichments but were not statistically significant. In addition, multiple histone modifications of epigenetic markers H3K9ac, H3K4me3 and H3K27ac, were all significantly enriched for cancer heritability. Repressed regions exhibited depletion (0.34-fold, p=1.2×10^−6^). Enrichment analysis of functional categories for each cancer subtype are shown in Supplementary Table 7.

Overall, cell-type-specific analysis of histone marks identified significant enrichments specific to individual cancers (Supplementary Fig. 2). For breast cancer, 3 out of 8 statistically significant tissues were adipose nuclei (H3K4me1, H3K9ac) and breast myoepithelial (H3K4me1) cells. For colorectal cancer, 15 out of the 18 statistical significant enrichments were observed in either colon or rectal tissues (colon/rectal mucosa, duodenum mucosa, small/large intestine and colon smooth muscle). We observed no significant enrichments for head/neck, lung and ovarian cancer, but we noted that for both lung (9 out of 10) and ovarian cancer (6 out of 10), the most enriched cell types were immune cells; while in head/neck cancer, 6 out of 10 most highly enriched cell types belonged to CNS (Supplementary Fig. 3, Supplementary Table 8). Cell-type-specific analysis for cancer subtypes are shown in Supplementary Table 9. Comparing cell-type-specific enrichment for cancers to the additional 38 non-cancer traits revealed notably differential clustering patterns (Supplementary Fig. 4). Breast, colorectal and prostate cancer showed enrichment mostly for adipose and epithelial tissues, in contrast to autoimmune diseases (enriched for immune/hematopoietic cells) or psychiatric disorders (enriched for brain tissues).

## DISCUSSION

We performed a comprehensive analysis quantifying the heritability and genetic correlation of six cancers, leveraging summary statistics from the largest cancer GWAS conducted to date. Our study demonstrates shared genetic components across multiple cancer types. These results contrast with a prior study conducted by Sampson *et al.* which reported an overall negligible genetic correlation among common solid tumors.^9^ Our results are, however, in line with a recent study^16^ which analyzed a subset of the data included here, and identified a significant genetic correlation between lung and colorectal cancer.

Our data support, and for the first time quantify, the strong genetic correlation (*r*_*g*_ =0.57) between lung and head/neck cancer, two cancers linked to tobacco use.^20^,^25^ We also for the first time observed a significant genetic correlation between breast and ovarian cancer (*r*_*g*_=0.24), two cancers that are known to share rare genetic factors including *BRCA1/2* mutations, and environmental exposures associated with endogenous and exogenous hormone exposures.^26^ Prostate cancer is also considered as hormone-dependent and associated with *BRCA1/2* mutations, but interestingly, we only observed a nominally significant and modest (*r*_*g*_=0.07) genetic correlation between breast and prostate cancer, whilst ovarian and prostate cancer showed no genetic correlation (*r*_*g*_=0.02, se=0.07).

Our large sample sizes allowed us to conduct well-powered analyses for cancer subtypes. While head/neck cancer showed negligible genetic correlation with overall (*r*_*g*_=0.03, se=0.06) and ER+ breast cancer (*r*_*g*_= −0.02, se=0.07), it showed a stronger genetic correlation with ER-breast cancer (*r*_*g*_ =0.21, se=0.09). Similarly, lung cancer showed a statistically more pronounced genetic correlation with ER-(*r*_*g*_=0.29, se=0.06) than ER+ breast cancer (*r*_*g*_=0.13, se=0.04). A recent pooled analysis of smoking and breast cancer risk demonstrated a smoking-related increased risk for ER+ but not for ER-breast cancer,^27^ and thus it is unlikely that the stronger genetic correlation between ER-subtype and lung and head/neck cancer is due to smoking behavior. Perhaps surprisingly, despite literature suggesting substantial similarities between ER-breast cancer and serous ovarian cancer in particular,^28^ we did not observe statistically significant different genetic correlations between ER-or ER+ breast cancer and serous ovarian cancer (*r*_*g*_ =0.17, se=0.08 *vs. r*_*g*_=0.11, se=0.06). This suggests that rare high penetrance variants may play a more important role in driving the similarities behind ER-breast cancer and serous ovarian cancer than common genetic variation.

Heritability analysis confirms that common cancers have a polygenic component that involves a large number of variants. Although susceptibility variants identified at genome-wide significance explain an appreciable fraction of the heritability for some cancers, we estimate that the majority of the polygenic effect is attributable to other, yet undiscovered variants, presumably with effects that are too weak to have been identified with current sample sizes. We found the genetic component that could be attributed to genome-wide significant loci varied greatly from ~0% for head/neck cancer to ~50% for breast and prostate cancer. These results reflect in part the strong correlation between number of GWAS-identified loci and sample size, as we had more than twice as many breast and prostate cancer samples compared to the other cancers. One corollary is that larger GWAS are likely to identify new susceptibility loci that could help our understanding of disease development, improve prediction power of genetic risk scores and hence contribute to screening and personalized risk prediction.^29^

Among the genetic correlations between cancer and non-cancer traits, we observed positive correlations for psychiatric disorders (depressive symptoms, schizophrenia) with lung and breast cancer, where findings from epidemiological studies have been suggestive but inconclusive. It has been proposed that the linkage between psychiatric traits and cancers are more likely to be mediated through cancer-associated risk phenotypes such as smoking, excessive alcohol consumption in depressed populations,^30^ and reduced fertility patterns (e.g., nulliparous) in psychiatric populations. Detailed analyses considering confounding traits like reproductive history and smoking are needed to make inference about the mechanisms involved. GWAS have identified pleiotropic regions influencing both lung cancer and nicotine dependence, such as 15q25.1.^32^,^33^ In line with those results, we identified a strong genetic correlation between smoking and both lung (*r*_*g*_=0.56) and head/neck cancer (*r*_*g*_=0.47). It remains unclear whether this genetic correlation is completely explained by the direct influence of smoking or if the shared genetic component affects the traits through separate pathways. Interestingly, a genetic correlation (*r*_*g*_=0.35, se=0.14) between lung and bladder cancer, another smoking-associated cancer, has been identified previously.^9^ Due to the small numbers of GWAS-identified smoking-associated SNPs, we were unable to assess a directional correlation between smoking and cancer, but we expect such analyses to become feasible as additional smoking-related SNPs are identified. We found modest positive, yet significant genetic correlations between adiposity-related measures (as reflected by waist-to-hip ratio, circulating HDL levels and BMI) and both colorectal and lung cancer, but negative genetic correlations between BMI and prostate and breast cancer, consistent with previous reported findings and reinforce the complex dynamics between obesity and cancer where multiple factors including age, smoking, endogenous hormones and reproductive status play a role.

We did not observe genetic correlations between breast cancer and age at menarche or age at natural menopause. These null observations were largely driven by ER+ breast cancer (ER+: *r*_*g*_= 0.006, se=0.03 *vs.* ER-: *r*_*g*_= −0.09, se=0.04 for age at menarche. ER+: *r*_*g*_= 0.0005, se=0.04 *vs.* ER-: *r*_*g*_= −0.10, se=0.05 for age at natural menopause), and were unexpected given that both factors play pivotal roles in breast cancer etiology and previous Mendelian randomization (MR) analyses have identified a link.^36,37^ An important difference between genetic correlation and MR analyses is that the latter only considers genome-wide significant SNPs while the former incorporates the entire genome. It is possible that a relatively small overlap in strongly associated SNPs can result in significant MR results despite low evidence of an overall genetic correlation. Indeed, the directional genetic correlations we observed for age at natural menopause, schizophrenia and HDL with breast cancer, and for lupus with prostate cancer, highlight again that although an overall genetic correlation may be negligible, there can still be genetic links between traits. It is important to note that we cannot rule out unmeasured confounding, including the possibility that these genetic variants affect an intermediate phenotype that is pleiotropic for both target traits. Given the observational nature of our data, these putative causal directions should be interpreted with caution.

Pan-cancer tumor-based studies have demonstrated that different cancers are sometimes driven by similar somatic functional events such as specific copy number abnormalities and mutations.^38^,^39^ Our enrichment results of germline genetic across functional annotation data shed new light on the biological mechanisms leading to cancer development. The more pronounced enrichment identified for conserved regions compared with coding regions provides evidence for the biological importance of the former, which has been shown to be true for multiple traits.^14^,^40^ Even though the biochemical function of many conserved regions remains uncharacterized, transcribed ultra-conserved regions have been found to be frequently located at fragile sites. Compared to normal cells, cancer cells have a unique spectrum of transcribed ultra-conservative regions, suggesting that variation in expression of these regions are involved in the malignant process.^41^,^42^ These results bridge the link between germline and somatic genetics in cancer development, which was also observed in a recent breast cancer GWAS that has demonstrated a strong overlap between target genes for GWAS hits and somatic driver genes in breast tumors.^43^ We also found a four-fold enrichment for transcription factor binding sites and a three-fold enrichment for super enhancers, consistent with prior observations that breast cancer GWAS loci fall in enhancer regions involved in distal regulation of target genes.^43^ Cell type-specific analysis of histone marks demonstrated the importance of tissue specificity, primarily for colorectal and breast cancer. Further, our results suggest that immune cells are important for ovarian and lung cancer whilst CNS is important to head/neck cancer. Unfortunately, we did not have data on prostate-specific tissues but we note that tissue-specific enrichment of prostate cancer heritability for epigenetic markers has been observed previously.^10^ We note that generation of rich functional annotation is ongoing and we expect to include additional tissue-specific functional elements in our future work.

Our study has several strengths. We were able to robustly quantify pair-wise genetic correlations between multiple cancers using the largest available cancer GWAS, comprising almost 600,000 samples across six major cancers and subtypes. We were also able to systematically assess the genetic correlations between cancer and 38 non-cancer traits. Notwithstanding the large sample sizes, several limitations need to be acknowledged. We did not have the sample sizes required to assess relevant cancer subgroups including oropharyngeal cancer, clear cell, mucinous and endometrioid ovarian cancer, or lung cancer among never smokers (each with ~2,000 cases). In addition, we did not have access to GWAS summary statistics for pre-*vs.* post-menopausal breast cancer. We were not able to consider all cancer risk factors when selecting non-cancer traits, since some of the well-established risk factors such as infection were either not available, showed no evidence of heritability or were not based on adequate sample sizes for robust analyses. SNP-heritability varies with minor allele frequency, linkage disequilibrium and genotype certainty; we note that approaches to estimate heritability leveraging GWAS data are constantly evolving. We also note that estimate variability needs to be taken into account when comparing the SNP-heritability with the classical twin-heritability, in particular for cancers with small sample sizes such as head/neck cancer (SNP-heritability varied between 5-14% and twin-heritability varied between 0-60%, although both point estimates were 9%). Further, our data were based on GWAS meta-analysis from multiple individual GWAS across European ancestry populations from Europe, Australia and the US. Intra-European ancestry differences are likely to be a source of bias. However, since we limited our analysis to SNPs with MAF>1% and HapMap3 SNPs (which have proven to be well-imputed across European ancestry populations), we believe that any population structure across cancers will have minimal effect on our results. Finally, as more non-European and multi-ethnic GWAS data become available, it is important to examine trans-ethnic genetic correlation in cancer.

In conclusion, results from our comprehensive analysis of heritability and genetic correlations across six cancer types indicate that solid tumors arising from different tissues share common germline genetic influences. Our results also demonstrate evidence for common genetic risk sharing between cancers and smoking, psychiatric and metabolic traits. In addition, functional components of the genome, particularly conserved and regulatory regions, are significant contributors to cancer heritability across multiple cancer types. Our results provide a basis and direction for future cross-cancer studies aiming to further explore the biological mechanisms underlying cancer development.

## METHODS

### Studies and quality control

We used summary statistics from six cancer GWASs based on a total of 597,534 participants of European ancestry. Cancer-specific sample sizes were: breast cancer: 122,977 cases / 105,974 controls; colorectal cancer: 36,948 / 30,864; head/neck cancer (oral and oropharyngeal cancers): 5,452 / 5,984; lung cancer: 29,266 / 56,450; ovarian cancer: 22,406 / 40,941; prostate cancer: 79,166 / 61,106. These data were generated through the joint efforts of multiple consortia. Details on study characteristics and subjects contributed to each cancer-specific GWAS summary dataset have been described elsewhere.^43-49^ SNPs were imputed to the 1000 Genomes Project reference panel (1KGP) using a standardized protocol for all cancer types.^18^ We included autosomal SNPs with a minor allele frequency (MAF) larger than 1% and present in HapMap3 (N_SNPs_ = ~1 million) because those SNPs are usually well imputed in most studies (note that excluding sex chromosomes could reduce the overall heritability estimates). A brief overview of the quality control in each cancer dataset are presented in Supplementary Table 10. For some of the cancers, we further obtained summary statistics data on subtypes (ER+ and ER-breast cancer; lung adenocarcinoma and squamous cell carcinoma; serous invasive ovarian cancer and advanced stage prostate cancer, defined as metastatic disease or Gleason score≥8 or PSA>100 or prostate cancer death). Sample sizes and more details shown in Supplementary Table 1.

We additionally assembled European ancestry GWAS summary statistics from 38 traits, which spanned a wide range of phenotypes including anthropometric (e.g., height and body mass index (BMI)), psychiatric disorder (e.g., depressive symptoms and schizophrenia) and autoimmune disease (e.g., rheumatoid arthritis and celiac disease) (Supplementary Table 11). We calculated trait-specific SNP-heritability and restricted our analysis to traits with a heritable component (Supplementary Table 12) as previously proposed.^14^ We removed the major histocompatibility complex (MHC) region from all analysis because of its unusual LD and genetic architecture.

### Estimation of SNP-heritability and genetic correlation

We estimated the SNP-heritability due to genotyped and imputed SNPs (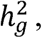, the proportion of phenotypic variance causally explained by common SNPs) of each cancer using LDSC.^15^ Briefly, this method is based on the relationship between LD score and χ^2^-statistics:

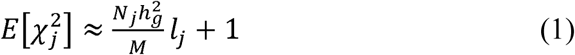

where 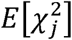 denotes the expected χ^2^-statistics for the association between the outcome and SNP*j*, N_j_ is the study sample size available for SNP*j*, M is the total numbers of variants and *l*_*j*_ denotes the LD score of SNP *j* defined as *l*_*j*_ = Σ_k_ *r*^2^(*j, k*) (k denotes other variants within the LD region). Note that the quantity estimated by LDSC is the causal heritability of common SNPs, which is different from the SNP-heritability as defined in Yang *et al.* To estimate 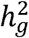 attributable to undiscovered loci, we identified SNPs that were associated with a given cancer at genome-wide significance (p<5×10-) and removed all SNPs +/-500,000 base-pairs of those loci prior to calculation (number of regions (+/-500 kb) for each cancer that reach the 5×10^−8^ threshold and measures of effect size are shown in Supplementary Table 2). We also converted the SNP-heritability from observed scale to liability scale by incorporating sample prevalence (*P)* and population prevalence (*F)* of each cancer:

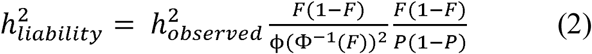

We subsequently calculated the genome-wide genetic correlations (*r*_*g*_) between different cancers, and between cancers and non-cancer traits, using an algorithm as previously described:^14^

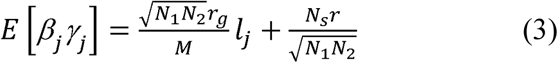

where *β*_*j*_ and *γ*_*j*_ are the effect sizes of SNP j on traits 1 and 2, *r*_*g*_ is the genetic covariance, M is number of SNPs, N_1_ and N_2_ are the sample sizes for trait 1 and 2, N_s_ is the number of overlapping samples, r is the phenotypic correlation in overlapping samples and *l*_*j*_ is the LD score defined as above. For genetic correlation between 6 cancers, the significance level is 0.05/15 = 0.003; for genetic correlation between 6 cancers and 38 traits, the significance level is 0.05/(6×38) = 0.0002.

Overall genetic correlations as estimated by LDSC are based on aggregated information across all variants in the genome. It is possible that even though two traits show negligible overall genetic correlation, there are specific regions in the genome that contribute to both traits. We therefore examined *local* genetic correlations between cancer pairs using ρ-HESS, an algorithm which partitions the whole genome into 1,703 regions based on LD-pattern of European populations and quantifies correlation between pairs of traits due to genetic variation restricted to these genomic regions. Local genetic correlation was considered statistically significant if p<0.05/1,703=2.9×10^−5^. In particular, we assessed the local genetic correlations for previously reported pleiotropic regions^18^,^51^ known to harbor SNPs affecting multiple cancers.

### Directional genetic correlation analysis

In addition to the genetic correlation analysis which reflects overall genetic overlaps, we also attempted to identify directions of potential genetic correlations using a subset of SNPs as proposed by Pickrell *et al.* The method adopts the following assumption: if a trait X influences trait Y, then SNPs influencing X should also influence Y, and the SNP-specific effect sizes for the two traits should be correlated. Further, since Y does not influence X, but could be influenced by mechanisms independent of X, genetic variants that influence Y do not necessarily influence X. Based on this assumption, the method proposes two “causal” models and two “non-causal” models, and calculates the relative likelihood ratio (LR) of the best non-causal model compared to the best causal model. We determined significant SNPs for each given cancer or trait in two independent ways, 1) LD pruned SNPs: we selected genome-wide significant (p<5×10^−8^) SNPs and pruned on LD-pattern in the European populations in Phase1 of 1KGP; 2) posterior probability of association (PPA) SNPs: we used a method implemented in “fgwas”^53^, which splits the genome into independent blocks based on LD-patterns in 1KGP and estimates the prior probability that any block contains an association. The model outputs posterior probability that the region contains a variant that influences the trait. We selected the lead SNP from each of the regions with a PPA of at least 0.9. We scanned through all pairs of cancers and traits to identify directional correlations. Only pairs of traits with evidence of directional correlations (LR comparing the best non-causal model over the best causal model<0.05) and without evidence of heteroscedasticity (pleiotropic effects) were reported as relatively more likely to exhibit mediated causation.

### Functional partitioning of SNP-heritability

To assess the importance of specific functional annotations in SNP-heritability across cancers, we partitioned the cancer-specific heritability using stratified-LDSC. This method partitions SNPs into functional categories and calculates category-specific enrichments based on the assumption that a category of SNPs is enriched for heritability if SNPs with high LD to that category have higher χ^2^ statistics than SNPs with low LD to that category. The analysis was performed using two previously described models.^14,24^

1) A full baseline-LD model including 24 publicly available annotations that are not specific to any cell type. When performing this model, we adjusted for MAF via MAF-stratified quantile-normalized LD score, and other LD-related annotations such as predicted allele age and recombination rate, as implemented by Gazal *et al*.^24^ Briefly, the 24 annotations included coding, 3’UTR and 5’UTR, promoter and intronic regions, obtained from UCSC Genome Browser and post-processed by Gusev *et al.*;^55^ the histone marks mono-methylation (H3K4me1) and tri-methylation of histone H3 at lysine 4 (H3K4me3), acetylation of histone H3 at lysine 9 (H3K9ac) processed by Trynka *et al.*^56-58^ and two versions of acetylation of histone H3 at lysine 27 (H3K27ac, one version processed by Hnisz *et al.*,^59^ another used by the Psychiatric Genomics Consortium (PGC)^60^); open chromatin, as reflected by DNase I hypersensitivity sites (DHSs and fetal DHSs),^55^ obtained as a combination of ENCODE and Roadmap Epigenomics data, processed by Trynka *et al.*;^58^ combined chromHMM and Segway predictions obtained from Hoffman *et al.*,^61^ which make use of many annotations to produce a single partition of the genome into seven underlying chromatin states (The CCCTC-binding factor (CTCF), promoter-flanking, transcribed, transcription start site (TSS), strong enhancer, weak enhancer categories, and the repressed category); regions that are conserved in mammals, obtained from Lindblad-Toh *et al.*^40^ and post-processed by Ward and Kellis;^62^ super-enhancers, which are large clusters of highly active enhancers, obtained from Hnisz *et al.*;^59^ FANTOM5 enhancers with balanced bi-directional capped transcripts identified using cap analysis of gene expression in the FANTOM5 panel of samples, obtained from Andersson *et al.*;^63^ digital genomic footprint (DGF) and transcription factor binding site (TFBS) annotations obtained from ENCODE and post-processed by Gusev *et al.*^55^

2) In addition to the baseline-LD model, we also performed analyses using 220 cell-type-specific annotations for the four histone marks H3K4me1, H3K4me3, H3K9ac and H3K27ac. Each cell-type-specific annotation corresponds to a histone mark in a single cell type (for example, H3K27ac in CD19 immune cells), and there were 220 such annotations in total. We further divided these 220 cell-type-specific annotations into 10 groups (adrenal and pancreas, central nervous system (CNS), cardiovascular, connective and bone, gastrointestinal, immune and hematopoietic, kidney, liver, skeletal muscle, and other) by taking a union of the cell-type-specific annotations within each group (for example, SNPs with any of the four histone modifications in any hematopoietic and immune cells were considered as one big category). When generating the cell-type-specific models, we added annotations individually to the baseline model, creating 220 separate models.

We performed a random-effects meta-analysis of the proportion of heritability over six cancers for each functional category. We set significance thresholds for individual annotations at p<0.05/24 for baseline model and at p<0.05/220 for cell-type-specific annotation.

## Data availability statement

The datasets generated during and/or analyzed during the current study are available from the authors on request.

Breast cancer: Summary results for all variants are available at http://bcac.ccge.medschl.cam.ac.uk/. Requests for further data should be made through the Data Access Coordination Committee (http://bcac.ccge.medschl.cam.ac.uk/).

Ovarian cancer: Summary results are available from the Ovarian Cancer Association Consortium (OCAC) (http://ocac.ccge.medschl.cam.ac.uk/). Requests for further data can be made to the Data Access Coordination Committee (http://cimba.ccge.medschl.cam.ac.uk/).

Prostate cancer: Summary results are publicly available at the PRACTICAL website (http://practical.icr.ac.uk/blog/).

Lung cancer: Genotype data for lung cancer are available at the database of Genotypes and Phenotypes (dbGaP) under accession phs001273.v1.p1. Readers interested in obtaining a copy of the original data can do so by completing the proposal request form at http://oncoarray.dartmouth.edu/

Head / neck cancer: Genotype data for the oral and pharyngeal OncoArray study have been deposited at the database of Genotypes and Phenotypes (dbGaP) under accession phs001202.v1.p1.

Colorectal cancer: Genotype data have been deposited at the database of Genotypes and Phenotypes (dbGaP) under accession number phs001415.v1.p1 and phs001078.v1.p1.

**Figure 5.**
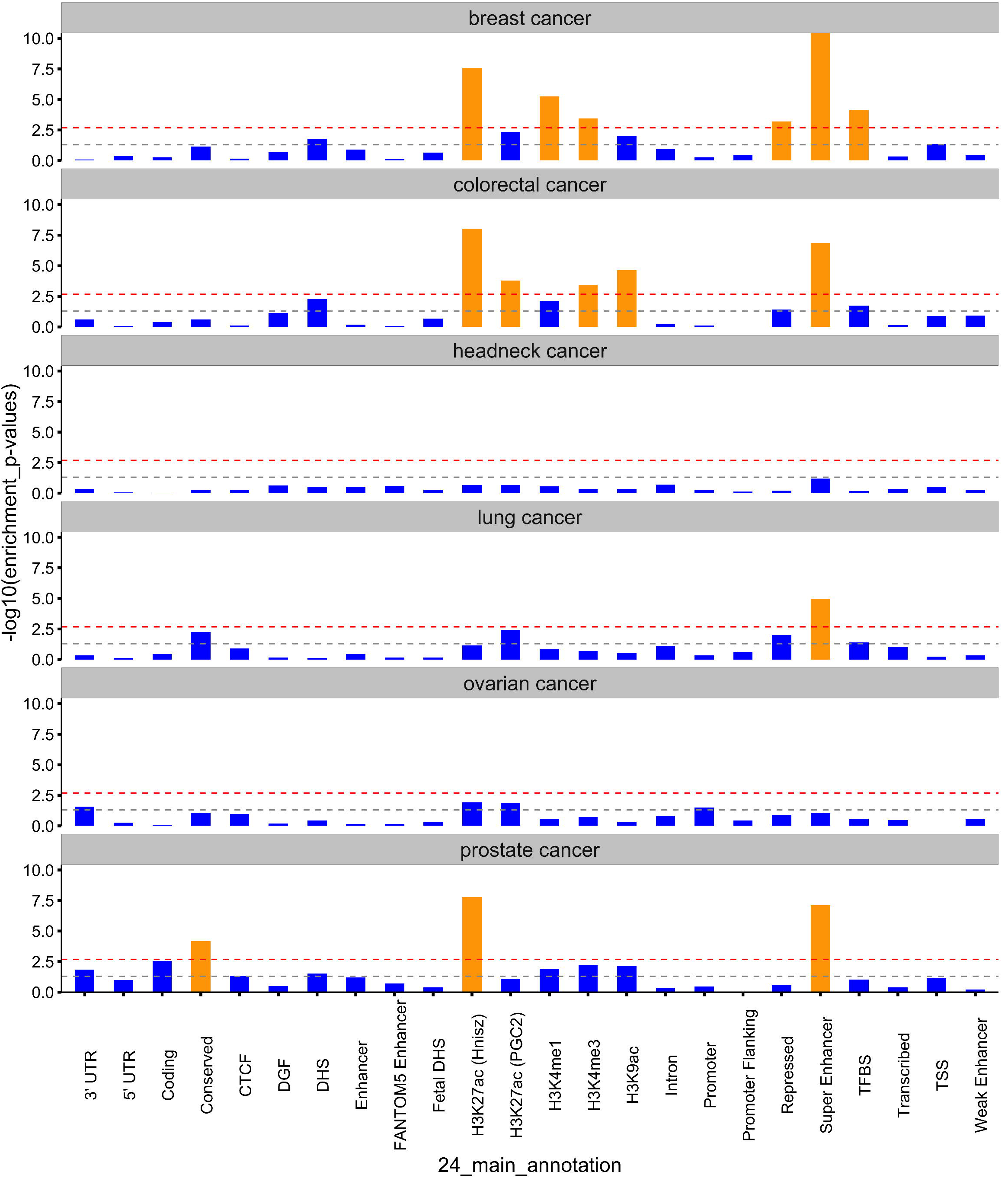
Enrichment p-values of 24 non-cell-type-specific functional categories over six cancer types. The x-axis represents each of the 24 functional categories, y-axis represents log-transformed p-values of enrichment. Annotations with statistical significance after Bonferroni corrections (p<0.05/24) were plotted in orange, otherwise blue. The horizontal grey dash line indicates p-threshold of 0.05; horizontal red dash line indicates p-threshold of 0.05/24. From top to bottom are six panels representing six cancers: breast cancer, colorectal cancer, head/neck cancer, lung cancer, ovarian cancer and prostate cancer. TSS: transcription start site; UTR: untranslated region; TFBS: transcription factor binding sites; DHS: DNase I hypersensitive sites; DGF: digital genomic foot printing; CTCF: CCCTC-binding factor.

## Author Contribution

All authors reviewed and commented on the manuscript, and approved the submission. C.I.A., C.L., D.F.E., F.R.S., P.J.B., P.D.P., P.K., S.L., S.L.S., S.B.G., X.J., R.J.H., K.M., M.M.G., and U.P. designed and managed the individual GWAS study; P.K., S.L., and X.J. developed and reviewed the analysis plan; S.L., X.J. analyzed the data with inputs from H.K.F; A.L.P., C.I.A., C.A.H., C.L., D.V.C., D.F.E., F.R.S., G.C., H.K.F., P.J.B., P.D.P., P.K., R.A.E., S.L., S.L.S., S.B.G., X.J., L.H., J.H., D.T., M.G., R.J.H., B.D., J.M., J.P.T., Y.H., K.M., K.B.K., J.D., M.M.G., J.S., B.P., and U.P. interpreted results; X.J. and S.L. drafted the manuscript with comments from A.L.P., C.I.A., C.A.H., C.L., D.V.C., D.F.E., F.R.S., G.C., H.K.F., P.J.B., P.D.P., P.K., R.A.E., S.L.S., S.B.G., L.H., J.H., D.T., M.G., R.J.H., B.D., J.M., J.P.T., Y.H., K.M., K.B.K., J.D., M.M.G., J.S., B.P., and U.P.

---

**Supplementary Figure 1.** QQ-plots showing region-specific p-values for the local genetic covariance for breast and colorectal cancer **(A)**, breast and head/neck cancer **(B)**, breast and lung cancer **(C)**, breast and ovarian cancer (**D**), breast and prostate cancer (**E**), colorectal and head/neck cancer (**F**), colorectal and lung cancer (**G)**, colorectal and ovarian cancer (**H**), colorectal and prostate cancer (**I**), head/neck and lung cancer (**J**), head/neck and ovarian cancer (**K**), head/neck and prostate cancer (**L**), lung and ovarian cancer (**M**), lung and prostate cancer (**N**), ovarian and prostate cancer (**O**). Each dot presents a specific genomic region. In the QQ plots, red color indicates significance after multiple corrections (p<0.05/1,703 regions compared), and blue color indicates nominal significance (p<0.05/15 pairs of cancers compared).

**Supplementary Figure 2. A)** Enrichment p-values of 220 cell-type-specific annotations in six major cancer types. The x-axis represents each of the 220 cell types, y-axis represents the log-transformed p-values of enrichment. Annotations with statistical significance after Bonferroni corrections (p<0.05/220) were plotted in orange, otherwise blue. The horizontal grey dash line indicates p-threshold of 0.05; horizontal red dash line indicates p-threshold of 0.05/220. The vertical green dash lines separate 220 cell types into ten cell type groups: adrenal and pancreas, cardiovascular, central nervous system, connective and bone, gastrointestinal, immune and hematopoietic system, kidney, liver, skeletal muscle, and others. From top to bottom are six panels representing six cancers: breast cancer, colorectal cancer, head/neck cancer, lung cancer, ovarian cancer, and prostate cancer. **B)** Enrichment p-values of the 220 cell-type-specific annotations meta-analyzed across six cancers.

**Supplementary Figure 3**. Enrichment of 220 cell-type-specific annotations in six major cancer types, plotted by histone marks (H3K4me1, H3K4me3, H3K9ac, H3K27ac). For each annotation, x-axis measures the proportion of SNPs accounted to that annotation, y-axis measures the proportion of heritability explained by that annotation. Annotations with statistical significance after Bonferroni corrections (p<0.05/220) are marked in red. Annotations with nominal significance (p<0.05) are marked in blue, the remaining annotations are marked in grey. **A)** breast cancer, **B)** colorectal cancer, **C)** head/neck cancer, **D)** lung cancer, **E)** ovarian cancer, and **F)** prostate cancer.

**Supplementary Figure 4.** Heat-maps showing bi-clustering of traits and cell-types over four histone marks. We performed 220 cell-type-specific annotation analysis in each of the 38 traits, and compared these enrichment results to the enrichment results of six cancers. Each checker reflects the beta coefficient z-score, scaled by traits. Red indicates enrichment, blue indicates depletion. Deeper color represents stronger magnitude of effects. The category of cell types is color coded to the left. **A)** H3K27ac, **B)** H3K4me1, **C)** H3K4me3 and **D)** H3K9ac. GI: gastrointestinal cell types; CNS: central nervous system cell types.

